# Multifaceted effects of bycatch mitigation measures on target/non-target species for pelagic longline fisheries and consideration for bycatch management

**DOI:** 10.1101/2022.07.14.500149

**Authors:** Daisuke Ochi, Kei Okamoto, Shintaro Ueno

## Abstract

The pelagic longline fishery, in an effort to reduce bycatch of sea turtles, have developed and deployed fisheries bycatch mitigation techniques such as replacing J/tuna hooks and squid bait with circle hooks and whole fish bait. However, little emphasis has been placed on the side effects of bycatch mitigation measures on endangered species other than target bycatch species. Several previous studies of the side effects have been marred by lack of control for the covariates. Here, based on long-term data obtained from research cruises by a pelagic longline vessel, we examined the effects of using circle hooks and whole fish bait to replace squid bait on the fishing mortality of target and non-target fishes, and also bycatch species. A quantitative evaluation analysis of our results, based on a Bayesian approach, showed the use of circle hooks to increase mouth hooking in target and bycatch species, and their size to be proportional to the magnitude of the effect. While deploying circle hooks did not increase fishing mortality per unit effort (MPUE) for shortfin mako sharks, combining to whole fish bait had a significant increase on MPUE. Because the impact of the introduction of bycatch mitigation measures on species other than the focused bycatch species is non-negligible, a quantitative assessment of bycatch mitigation-related fishing mortality is critical before introducing such measures.

## INTRODUCTION

Unintentional catch in fisheries is known as bycatch, and bycatch of particularly endangered species, can have devastating effects on these populations. Therefore, efforts to minimize bycatch of endangered species are strongly encouraged at all levels of conservation from local to international. For example, in the tuna longline fishery, concerns about the increased conservation risk for seabirds and sea turtles by unintentional and fatal catch have been a major issue among many regional fisheries management organizations (RFMOs) since the 1990s (Wallace et al. 2013; Dias et al. 2019). In recent years, some elasmobranchs, whose populations are declining further, are also treated as bycatch species. The decline of these species has focused attention at both the national and the international levels. Even for target fish that are not bycatch species, addressing the deterioration of stock status caused by overfishing requires reductions in unintentional fishing mortality (for instance, billfishes in the North Atlantic; Kerstetter & Graves 2006; Diaz 2008).

Several studies have weighed the sustainability of tuna longline fisheries against the conservation of species vulnerable to bycatch (Hall et al. 2000; Melvin et al. 2014; Clarke et al. 2015)—particularly for seabirds and marine turtles—with the development of several effective bycatch mitigation measures (Melvin et al. 2014; Swimmer et al. 2017). Bycatch mitigation measures in longline fisheries target specific animal groups and are evaluated based on their success in reducing mortality due to bycatch of specific vulnerable species. While the impact on the catch of the target fish is the primary consideration when evaluating bycatch mitigation techniques, few studies have examined the impacts and tradeoffs on species not targeted by mitigation techniques (Pacheco et al. 2011; Gilman et al. 2016). However, the introduction of bycatch mitigation measures requires an assessment of the impact not only on the target bycatch species on the other ecologically related species prior to their introduction (Reinhardt et al. 2018).

Use of circle hook and whole finfish bait are typical sea turtle bycatch mitigation measures for pelagic longline fisheries (Watson et al. 2005; Gilman et al. 2006; Yokota et al. 2009; Stokes et al. 2011). The tip of the circle hook bends inward, and when a fish or sea turtle swallows the hooked bait, the circle hook less likely to hook inside the digestive tract; instead, as the hook exits the mouth, a torque causes it to hook through the edge of the mouth. This property allows for easy hook removal and has been reported to reduce the mortality rate of bycatch sea turtles on board and after release (Kiyota et al. 2004; Cooke & Suski 2004; Kerstetter & Graves 2006). Reports of positive effects of circle hooks include those for other species—such as reduced haulback and post-release mortality in sharks, reduced post-release mortality in swordfish (*Xiphias gladius*), and increased catch rates in tuna, the target fish. The use of fish bait instead of squid bait potentially reduces bycatch (Watson et al. 2005; Yokota et al. 2009, 2011) and mortality rates (Stokes et al. 2011; Parga et al. 2015) of sea turtles. These sea turtle mitigation measures, however, exact other costs. Several meta-analytic studies have reported the circle hook related reduction of sea turtle mortality rate but increase in billfish and shark catch rates (Gilman et al. 2016; Reinhardt et al. 2018; Santos et al. 2020). The use of whole fish bait also reportedly increases the catch of sharks and other species (Foster et al. 2012). However, few studies have allowed for quantitative evaluation of the effects of these mitigation measures experimentally on species beyond sea turtles. This limited data concern is due in part to the reliance on observer data from commercial vessels, and small sample sizes, small comparison groups, and lack of experimental rigor from research vessels. In addition, the impact assessment for sharks underestimates catch rates and mortality associated with missed catches due to “bite-off” branchline (Reinhardt et al. 2018). In addition, many experimental studies and meta-analyses (e.g., Diaz 2008; Godin et al. 2012; Huang et al. 2016; Pacheco et al. 2011; Yokota et al. 2006a) only evaluate gear impacts at significance levels without evaluating the magnitude of the effect. Small significant differences may be judged not to matter much when assessing overall risk in bycatch species. Studies that controlled for these conditions would allow for an evaluation of adverse effects without the confounding problems described above. Also, many studies use catch/bycatch rates (Andraka et al. 2013; Foster et al. 2012; Gilman et al. 2007; Watson et al. 2005; Yokota et al. 2006a) and mortality rates (at haulback or after release; Carruthers et al. 2009; Gallagher et al. 2014; Horodysky et al. 2005; Kerstetter et al. 2003) as important impact indicators, without considering irreversible impacts, such as the number of organisms killed at the time of catch. Additionally, hooking location itself (mouth, swallow, or external) is believed to have a strong influence on mortality rate, which demands the estimation of risk under specific hooking conditions, taking causal relationships into account.

Here, we used data from controlled experiments to analyze these confounding factors and developed a Bayesian statistical model to evaluate the effects of changing hook and bait type on fish species other than turtles—particularly, tuna, swordfish, and sharks—and then verified the contribution of circle hooks and fish bait to mortality rate, catch rate, and fatal catch rate, respectively. We also discussed the appropriate assessment of bycatch mitigation measures in fisheries management.

## METHOD

### Experimental Operations

We analyzed data from the R/V Taikei No. 2 longline research operation conducted in the Northwest Pacific Ocean between 2002 and 2010—a typical Japanese shallow-setting operation targeting mainly swordfish and sharks (set depth: shallowest 47.44m±14.22SD, deepest 72.41m±10.87SD (n=98 ops.); based on the time depth recorder data [SBT500, Murayama Denki Ltd.]), using four hooks per basket, a wire leader, and a night soaking (Yokota et al. 2006a). A total of 286,363 hooks from 306 operations (range: 400 – 964 hooks per operation) were deployed in the experiment (Table 1). The area of operation ranged from around the Izu Islands in Japan to off the east coast of northeastern Honshu—typical fishing ground for Japanese shallow-setting longliners (Fig. 1; Hiraoka et al. 2016). We deployed 11 different hooks (Table 2, Appendix S1, S2) and we describe hook shapes and other details of these hooks in Appendix S1 following the measurement method of Yokota et al. (2006b). Because degree of hook offset has been reported to affect catch and hooking location (Cooke & Suski 2004), most hooks were <10° but some were nearly 15°. The bait comprised chub mackerel (*Scomber japonicus*) and Japanese common squid (*Todarodes pacificus*) in the range of 20–30 cm in fork length or dorsal mantle length. These were frozen and stored, then completely thawed before being hooked. The sequence of line setting was divided into several experimental segments, and a different combination of hook and bait type was applied for each segment, with an alternate order of segments at each operation.

**Figure 1.**
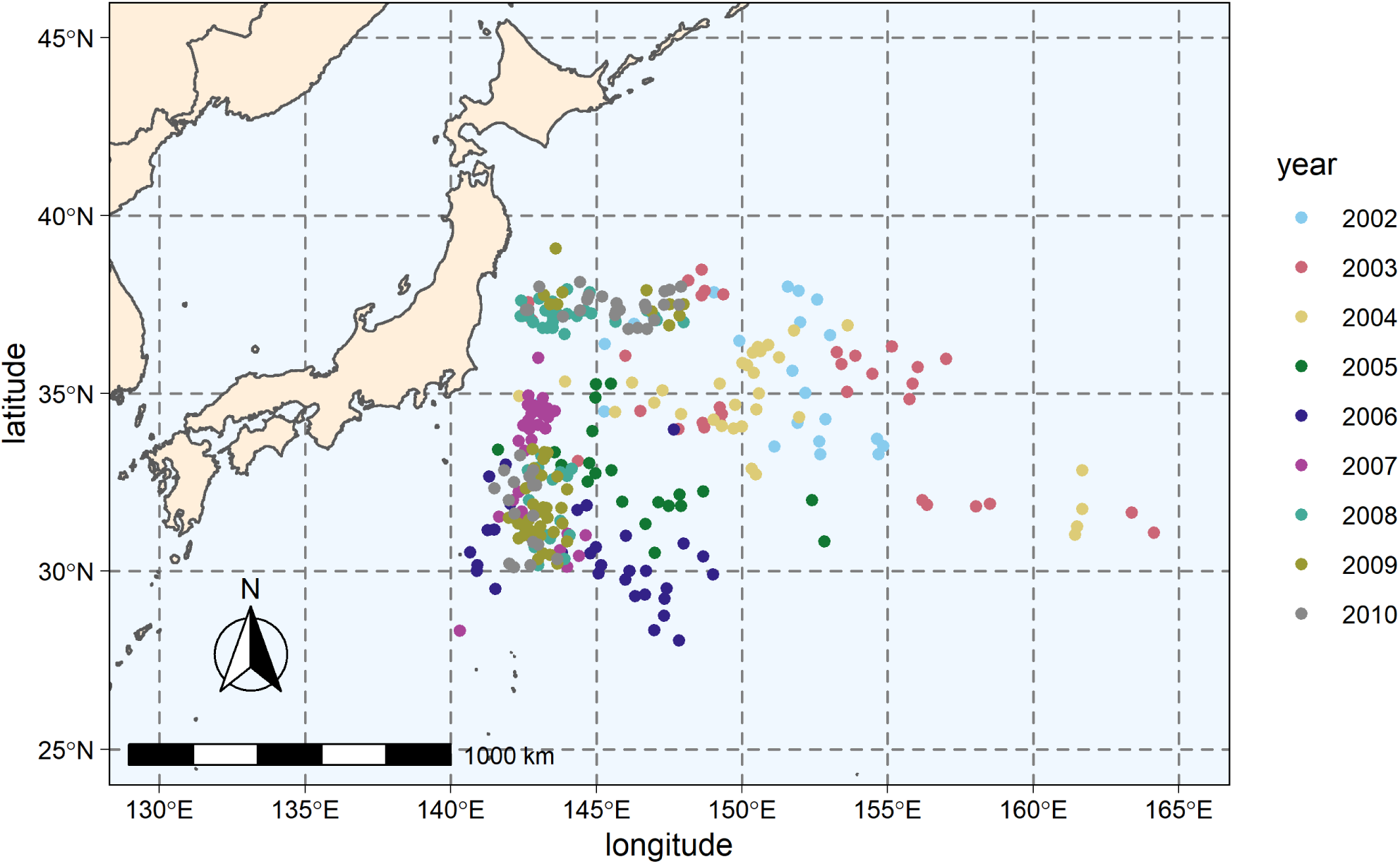
Locations where the longline operation experiment was conducted.

**Table 1.** Fishing effort (longline hooks) in experimental operations used in the analysis.

| Hook type | Bait type |  | Total effort |
| --- | --- | --- | --- |
|  | squid | fish |  |
| tuna | 97,834 | 71,146 | 168,980 |
| small-C | 70,882 | 7,658 | 78,540 |
| large-C | 31,872 | 6,976 | 38,848 |
| total effort | 200,588 | 85,780 | 286,368 |

**Table 2.** List of hook types used in the experiment.

| hook name | size | category | straight total<br>length(mm) | maximum total<br>width(mm) | minimum total<br>width(mm) |
| --- | --- | --- | --- | --- | --- |
| Komatsu Keisaku tunahook | 4.0 sun | tuna | 63 | 63 | 38 |
| Doitomi Tunamutsu | 4.0 sun | small-C<br>(circle hook) | 58 | 69 | 39 |
| Komatsu Keisaku modified | 4.0 sun | small-C<br>(circle hook) | 60 | 68 | 41 |
| Komatsu Keisaku modified | 4.5 sun | small-C<br>(circle hook) | 63 | 71 | 45 |
| Komatsu Keisaku modified | 4.8 sun | small-C<br>(circle hook) | N/A* | N/A* | N/A* |
| Komatsu Keisaku type Koshina | 4.5 sun | small-C<br>(circle hook) | 62 | 76 | 51 |
| Komatsu Keisaku type North America | 4.3 sun | small-C<br>(circle hook) | 57 | 63 | 41 |
| Tankichi Uruwa | 3.8 sun | small-C<br>(circle hook) | 56 | 64 | 36 |
| Komatsu Keisaku modified | 5.2 sun | large-C<br>(circle hook) | 74 | 81 | 48 |
| Komatsu Keisaku type North America | 5.2 sun | large-C<br>(circle hook) | 76 | 85 | 52 |
| Pacific Fishing Tackle circle hook | 18/0 | large-C<br>(circle hook) | 68 | 80 | 51 |

\* No measurement data were available due to the loss of the hook after the experiment.

The researcher recorded catch for each operation for all catch/bycatch, and the species caught, fate of catch (alive or dead), hooking location (mouth, swallow, and external hooking), time of catch, and float ID. The researchers determined if the catch was alive or dead based on the movement of the animals and the degree of injuries before being hauled. Float ID was recorded when a float was dropped during line setting and when it was retrieved to the deck during line hauling in order to calculate soak time. Hooking locations were recorded for catches caught using squid bait. At the start of longline operations, the researcher also collected sea surface temperature with a water thermometer (DS-1; Murayama Denki Ltd.) equipped on the vessel, which we subsequently used in the analysis.

The experiment was conducted using multiple sizes of circle hooks, which varied greatly in size. However, those sample sizes were too small to analyze each hook type separately. For convenience, we have classified the hook shapes based on size (threshold: straight total length = 68 mm AND maximum total width = 80 mm; approximately equivalent to 5.0-sun or 18/0). This led to the following three main hook types:

1. control (tuna hook 4.0-sun; tuna)
2. smaller circle hook (smaller than the threshold; small-C)
3. larger circle hook (the threshold or larger; large-C)

In Table 1, we show longline effort separated by bait and hook type, tabulated according to the above categorization. We selected the following species based on whether there was a sufficient sample size to statistical analyses (especially for model convergence): bigeye tuna (*Thunnus obesus*), blue shark, common dolphinfish (*Coryphaena hippurus*), escolar (*Lepidocybium flavobrunneum*), longnose lancetfish (*Alepisaurus ferox*), shortfin mako shark, striped marlin (*Kajikia audax*), swordfish, and loggerhead turtle (*Caretta caretta*).

### Statistical Analysis

We conducted all analyses using a Bayesian approach to estimate parameters. We adopted haulback mortality rate, catch per unit effort (CPUE; /1000 hooks), and mortality per unit effort (MPUE; /1000 hooks) (Afonso et al. 2011) as indices to evaluate the impact of hook and bait type on fishing mortality within the analyses. We used the following data as inputs to the model: number of caught (individuals), longline effort (number of hooks), fate at hauling, hooking location, year and location of operation, water temperature at operation, and soaking time.

Because of the missing data due to not recording the hooking location when fish bait was used, we split the analysis to evaluate the impact of hook type and bait type on fishing mortality into two models. MODEL 1 evaluated only the effect of hook type based on capture events with squid bait and assumed that the use of circle hooks would change to more mouth hooking of each species, resulting in improved mortality rate. MODEL 2 evaluated both hook and bait types, and assumed that the combination of the two would result in large fluctuations in mortality rate, CPUE and MPUE for each species.

We based MODEL 1 on a logit regression using a Bernoulli distribution. We express the observed hooking location *H* and haulback mortality rate *M* in MODEL 1 by the following equations:

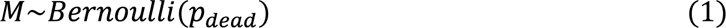

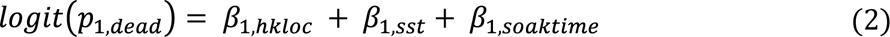

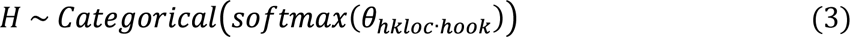

where *β_1_* is the parameter in each explanatory variable (*hkloc:* hooking location, *sst:* sea surface temperature and *soaktime:* soak time), *p_1,dead_* is the expected haulback mortality rate, and *θ_hklocꞏhook_* is the expected probability of hooking location in each hook type.

We structured MODEL 2 to calculate the expected number of mortalities per effort (MPUE) based on the parameters estimated in both the mortality rate and CPUE estimation subsets. In MODEL 2, due to lack of hooking location data, we calculate the mortality rate *p_2,dead_* from a modified equation (2) as follows;

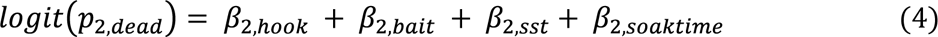

where *β_2_* is the parameter in each explanatory variable (*hook:* hook type, *bait:* bait type).

The CPUE subset is based on a log regression using a Poisson distribution as the error structure. We express the number of catches *C* per operation using the expected CPUE *λ* as follows:

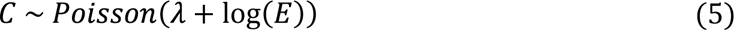

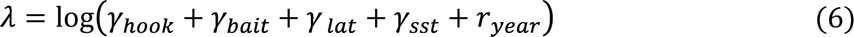

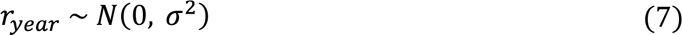

where *E* is the longline effort (hooks per set), *γ* is a parameter in each explanatory variable (*lat:* latitude where the longline set), *r_year_* is a random effect of annual fluctuation on CPUE, and *σ* is a standard deviation.

We obtained the expected values of hook-bait-specific MPUE ζ by multiplying CPUE by at-haulback mortality rate as follows:

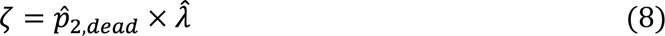

Where 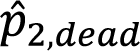 denotes the expected mortality rate standardized for hook and bait type, and 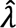 denotes the estimated CPUE standardized for hook and bait type. We used the standardization method for abundance indices used in fisheries stock assessment (Maunder & Punt 2004). In this method, explanatory variables other than the factor subject to standardization are averaged to predict the objective variable, which in stock assessment is a time scale such as years or months, but in our case, we modified the standardization scale to reference hook type and bait type.

We calculated each parameter based on a Bayesian approach with Markov chain Monte Carlo (MCMC) sampling. For the MCMC sampling, we used cmdstan 2.28.2 (Stan Development Team 2021). As a prior distribution for σ, we used a half student-t distribution with 2 degrees of freedom, mean 0, and variance 2.5, and we used a uniform distribution for the other parameters (*β_1_, β_2_, γ, θ*). We computed the posterior distribution using Stan with 15000 sampling iterations including 10000 warmup iterations, number of chains as 4, and no sinning. We calculated Bayesian credible intervals based on the highest density interval (HDI) for the estimates. Although the Bayesian approach for the estimates precluded significance testing, we determined the difference between the estimates of the experimental group and those of the control group (“swallowing” for hooking location, “tuna” hook for hook type, and “squid” for bait type), and if the lower and upper limits of HDI for the difference did not exceed 0, we considered the difference as a difference for convenience (assuming region of practical equivalence [ROPE] as 0; Kruschke 2015). In Appendix S3 we show the Stan code used to estimate each parameter in MODEL 1 and 2. For other data handling, statistical analysis, and plotting, we used R4.3.0 (R Core Team 2023) and packages “cmdstanr 0.6.1,” “ggalluvial 0.12.5,” “ggthemes 4.2.4,” “mapdata 2.3.1,” “maps 3.4.1,” “sf 1.0–14,” “tidybayes 3.0.6,” and “tidyverse 2.0.0” (Pebesma 2018; Wickham et al. 2019; Becker et al. 2020; Brunson 2020; Arnold 2021; Gabry & Češnovar 2021; Becker et al. 2022; Kay 2023).

## RESULTS

### Summary Statistics

Sufficient catches/bycatches of blue shark, longnose lancetfish, common dolphinfish, shortfin mako shark swordfish, bigeye tuna, loggerhead turtle, escolar and striped marlin were recorded for the later analysis (Table 3). The main species listed in Table 3 as “other species” are listed as follows—salmon shark (*Lamna ditropis*; N = 229), pelagic stingray (*Pteroplatytrygon violacea*; N = 199), pomflets (*Brama spp.*; N = 135), bigeye thresher shark (*Alopias superciliosus*; N = 89), and albacore (*Thunnus alalunga*; N = 69). The sample sizes of these “other species” were too skewed among experimental groups to converge the later analysis.

**Table 3.** Number of individuals caught by hook and bait type in the experimental operation.

| Species | Total<br>catch | Hook type |  |  | Bait type |  |
| --- | --- | --- | --- | --- | --- | --- |
|  |  | tuna | small-C | large-C | squid | fish |
| blue shark | 13,018 | 8,084 | 3,562 | 1,372 | 8,903 | 4,115 |
| longnose lancetfish | 1,297 | 945 | 257 | 95 | 692 | 605 |
| common dolphinfish | 505 | 206 | 181 | 118 | 363 | 142 |
| shortfin mako shark | 485 | 298 | 134 | 53 | 262 | 223 |
| swordfish | 288 | 129 | 112 | 47 | 249 | 39 |
| bigeye tuna | 269 | 114 | 146 | 9 | 201 | 68 |
| loggerhead turtle | 268 | 128 | 113 | 27 | 259 | 9 |
| escolar | 163 | 79 | 56 | 28 | 108 | 55 |
| striped marlin | 145 | 70 | 53 | 22 | 126 | 19 |
| other species | 1,578 | - | - | - | - | - |

In Table 3, we also show the number of fish caught by hook type and bait type. Bigeye tuna had extremely low catches on large-C hook, and loggerheads had low catches on fish bait. The most common hooking location at the time of catch was mouth hooking for all nine species, with extremely few hook locations other than mouth hooking, especially for bigeye tuna and escolar (Table 4). The proportion of mortality of captured species at haulback varied greatly by species. The haulback mortality rate was low for blue shark, common dolphinfish, escolar and shortfin mako shark, and higher for bigeye tuna, longnose lancetfish, striped marlin and swordfish. In the case of loggerhead turtles, the mortality rate was extremely low.

**Table 4.** Composition of hooking location and fate at hauling.

| Species | Hooking location |  |  |  | Fate at haulback |  |  |
| --- | --- | --- | --- | --- | --- | --- | --- |
|  | swallowed | mouth | external | unknown (*) | alive | dead | unknown (*) |
| blue shark | 2,270 | 2,608 | 66 | 8,074 | 11,701 | 1,066 | 251 |
| longnose lancetfish | 25 | 288 | 16 | 968 | 165 | 1,010 | 122 |
| common dolphinfish | 39 | 159 | 4 | 303 | 398 | 87 | 20 |
| shortfin mako shark | 66 | 68 | 18 | 333 | 363 | 118 | 4 |
| swordfish | 59 | 118 | 18 | 93 | 55 | 226 | 7 |
| bigeye tuna | 7 | 128 | 2 | 132 | 83 | 183 | 3 |
| loggerhead turtle | 109 | 116 | 14 | 29 | 256 | 5 | 7 |
| escolar | 1 | 62 | 1 | 99 | 108 | 40 | 15 |
| striped marlin | 12 | 80 | 8 | 45 | 68 | 76 | 1 |

(\*) Includes catches that dropped off before reseachers checked or lack of survey.

The length-based size composition of the nine species (precaudal length for blue sharks and shortfin mako sharks, straight-line carapace length for loggerhead turtles, eye-to-fork length for striped marlin and swordfish, and fork length for all other species) used in the analysis was not statistically compared in this study and was not included in the model because it did not contribute to haulback mortality rate, but in Appendix S4 we included histograms.

Almost all parameters in the models for the nine species were successfully converged (Rhat < 1.1) but the whole models were not converged for bigeye tuna, common dolphinfish, escolar in the MODEL 1, and some of the parameters in the MOEDL 1 and the whole model in the MODEL 2 for loggerhead turtle were not converged, as we describe below.

### Output of MODEL 1

In Table 5 we show occurrence estimates of hooking locations by hook type throughout the model. We observed differences in hooking location by hook type with an increase in mouth hooking and a decrease in hook swallowing for large-C for loggerhead turtle and a clear increase in mouth hooking for small-C for shortfin mako shark and swordfish (Fig. 2). In blue sharks, the frequency of hook swallowing decreased in both small-C and large-C.

**Figure 2.**
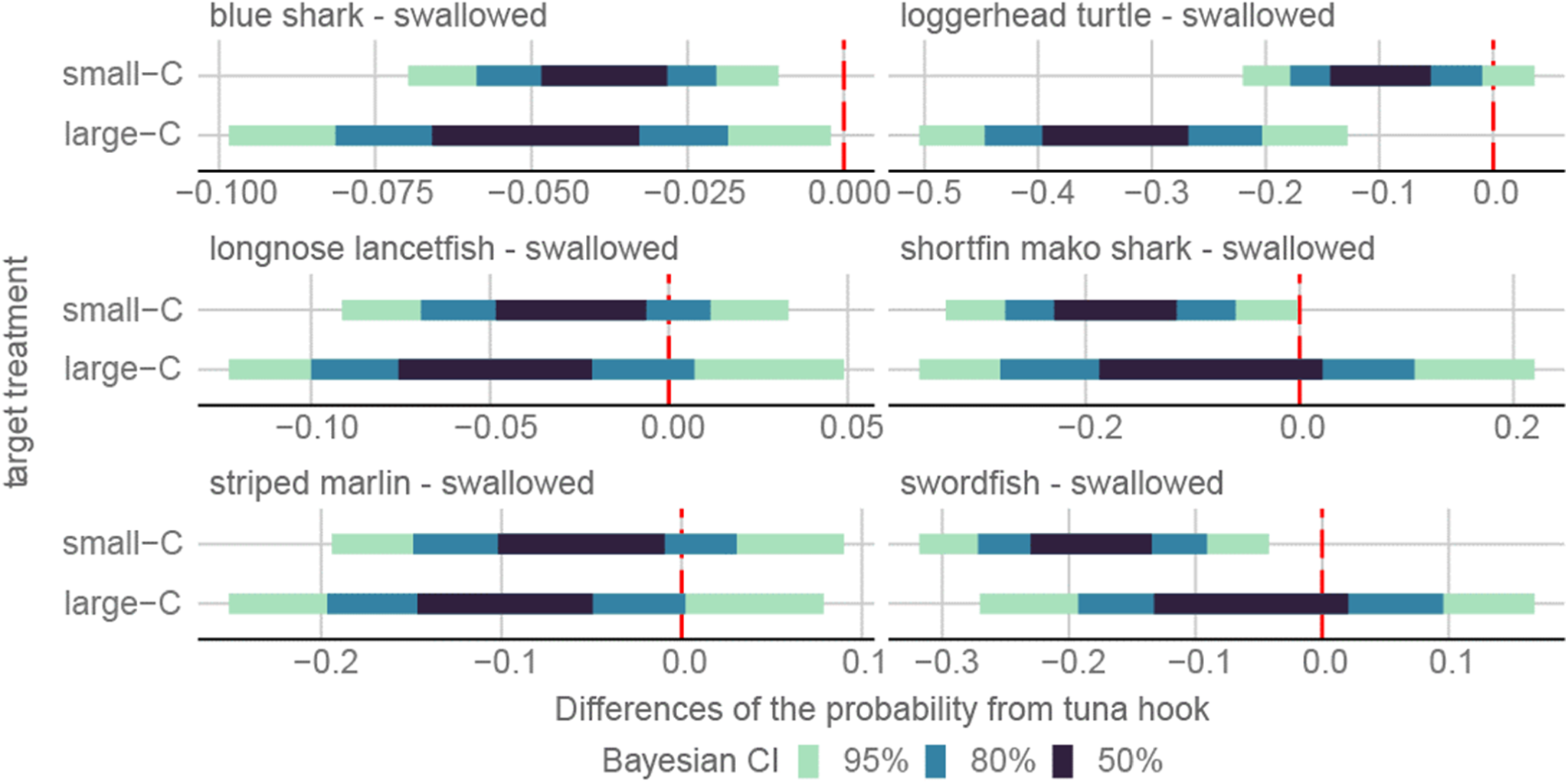
Differences in the estimated probability of the “swallowed” hooking location of each circle hook type from tuna hook when squid bait is used. The red dotted line indicates that the difference is zero.

**Table 5.**
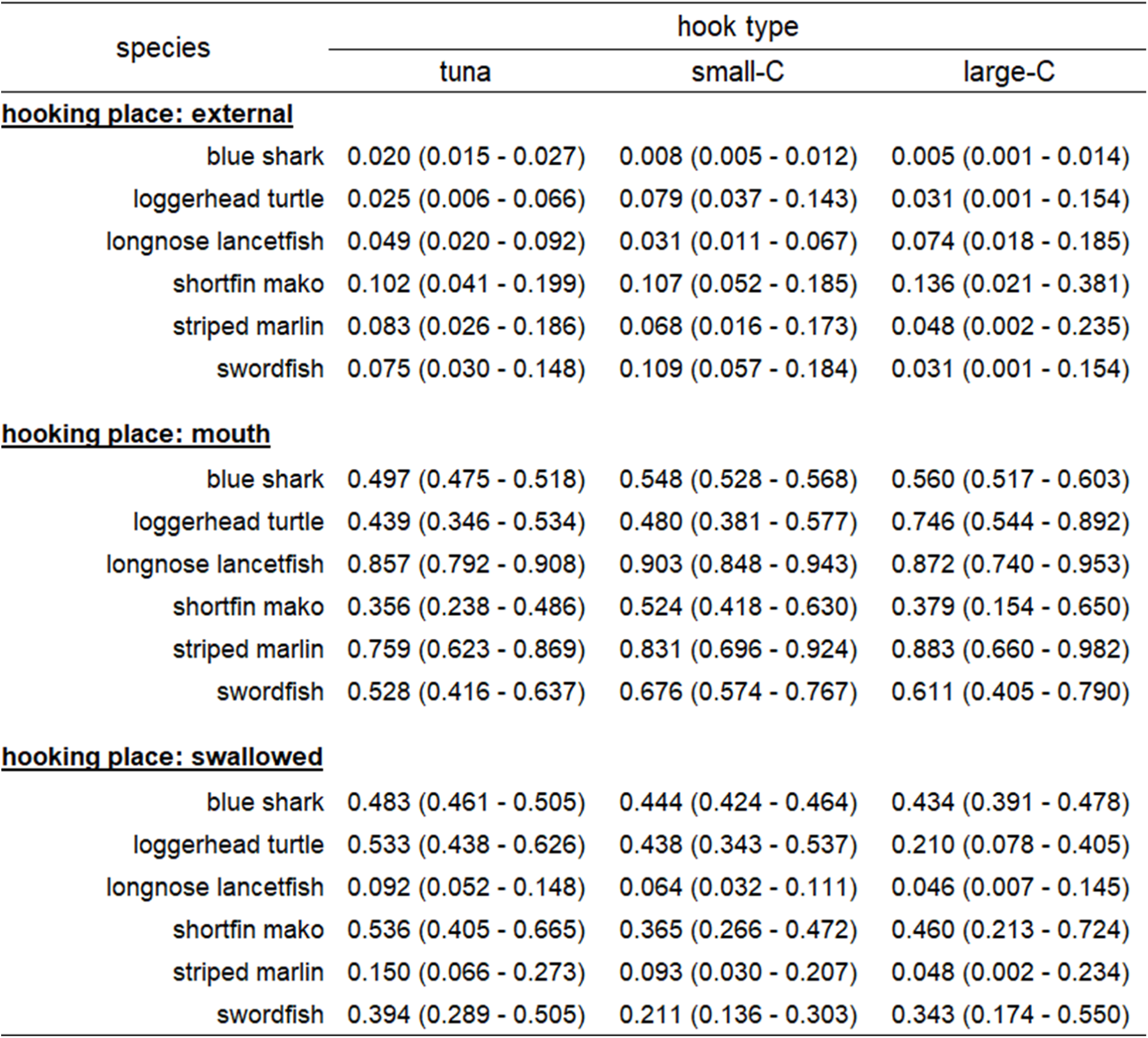
Estimates of the posterior distribution of the proportion of hooking location by hook when squid bait is used (median). Lower and upper limits of Bayesian credible interval (95% highest density interval [HDI]) are shown in parentheses.

| species | hook type |  |  |
| --- | --- | --- | --- |
|  | tuna | small-C | large-C |
| **<u>hooking place: external</u>** |  |  |  |
| blue shark | 0.020 (0.015 - 0.027) | 0.008 (0.005 - 0.012) | 0.005 (0.001 - 0.014) |
| loggerhead turtle | 0.025 (0.006 - 0.066) | 0.079 (0.037 - 0.143) | 0.031 (0.001 - 0.154) |
| longnose lancetfish | 0.049 (0.020 - 0.092) | 0.031 (0.011 - 0.067) | 0.074 (0.018 - 0.185) |
| shortfin mako | 0.102 (0.041 - 0.199) | 0.107 (0.052 - 0.185) | 0.136 (0.021 - 0.381) |
| striped marlin | 0.083 (0.026 - 0.186) | 0.068 (0.016 - 0.173) | 0.048 (0.002 - 0.235) |
| swordfish | 0.075 (0.030 - 0.148) | 0.109 (0.057 - 0.184) | 0.031 (0.001 - 0.154) |
| **<u>hooking place: mouth</u>** |  |  |  |
| blue shark | 0.497 (0.475 - 0.518) | 0.548 (0.528 - 0.568) | 0.560 (0.517 - 0.603) |
| loggerhead turtle | 0.439 (0.346 - 0.534) | 0.480 (0.381 - 0.577) | 0.746 (0.544 - 0.892) |
| longnose lancetfish | 0.857 (0.792 - 0.908) | 0.903 (0.848 - 0.943) | 0.872 (0.740 - 0.953) |
| shortfin mako | 0.356 (0.238 - 0.486) | 0.524 (0.418 - 0.630) | 0.379 (0.154 - 0.650) |
| striped marlin | 0.759 (0.623 - 0.869) | 0.831 (0.696 - 0.924) | 0.883 (0.660 - 0.982) |
| swordfish | 0.528 (0.416 - 0.637) | 0.676 (0.574 - 0.767) | 0.611 (0.405 - 0.790) |
| **<u>hooking place: swallowed</u>** |  |  |  |
| blue shark | 0.483 (0.461 - 0.505) | 0.444 (0.424 - 0.464) | 0.434 (0.391 - 0.478) |
| loggerhead turtle | 0.533 (0.438 - 0.626) | 0.438 (0.343 - 0.537) | 0.210 (0.078 - 0.405) |
| longnose lancetfish | 0.092 (0.052 - 0.148) | 0.064 (0.032 - 0.111) | 0.046 (0.007 - 0.145) |
| shortfin mako | 0.536 (0.405 - 0.665) | 0.365 (0.266 - 0.472) | 0.460 (0.213 - 0.724) |
| striped marlin | 0.150 (0.066 - 0.273) | 0.093 (0.030 - 0.207) | 0.048 (0.002 - 0.234) |
| swordfish | 0.394 (0.289 - 0.505) | 0.211 (0.136 - 0.303) | 0.343 (0.174 - 0.550) |

In Table 6 we show haulback mortality rates by hooking location. We observed clear differences in haulback mortality rate by hooking location for blue shark, shortfin mako shark, striped marlin, and swordfish (Fig. 3). Haulback mortality rate after mouth hooking for blue sharks was lower, and that of external hooking was higher than those for hook swallowing. We observed lower haulback mortality rates for shortfin mako shark, striped marlin, and swordfish from mouth hooking than from hook swallowing.

**Figure 3.**
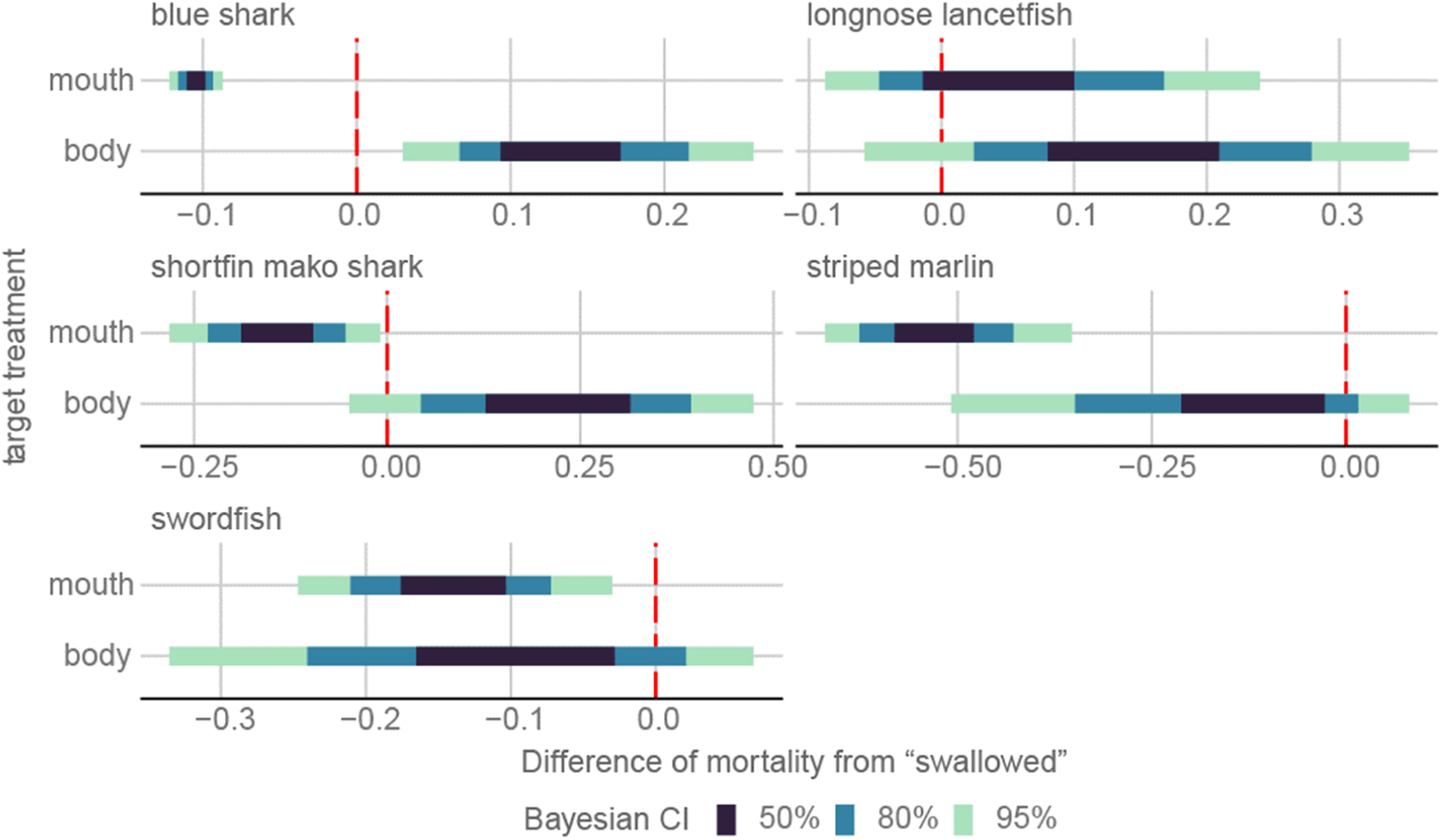
Differences in the estimated haulback mortality rate of each target hooking location from “swallowed” hooking location when squid bait is used. The red dotted line indicates that the difference is zero.

**Table 6.**
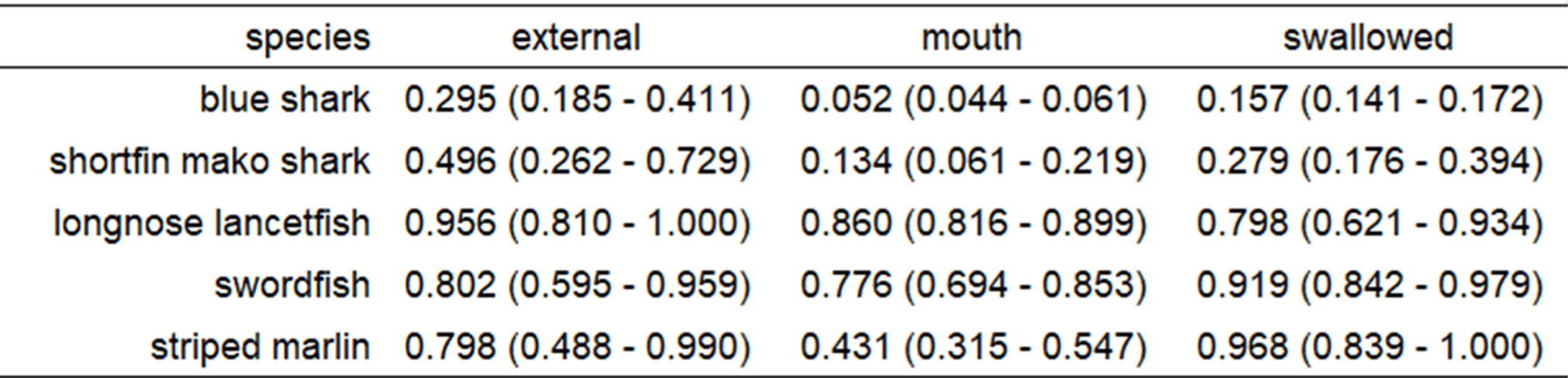
Estimated haulback mortality rates (median of posterior distribution) by hooking location when squid bait is used. Lower and upper limits of Bayesian credible interval (95% highest density interval [HDI]) are shown in parentheses. Loggerhead turtles were excluded because there were no mortalities and the calculation had not been converged.

| species | external | mouth | swallowed |
| --- | --- | --- | --- |
| blue shark | 0.295 (0.185 - 0.411) | 0.052 (0.044 - 0.061) | 0.157 (0.141 - 0.172) |
| shortfin mako shark | 0.496 (0.262 - 0.729) | 0.134 (0.061 - 0.219) | 0.279 (0.176 - 0.394) |
| longnose lancetfish | 0.956 (0.810 - 1.000) | 0.860 (0.816 - 0.899) | 0.798 (0.621 - 0.934) |
| swordfish | 0.802 (0.595 - 0.959) | 0.776 (0.694 - 0.853) | 0.919 (0.842 - 0.979) |
| striped marlin | 0.798 (0.488 - 0.990) | 0.431 (0.315 - 0.547) | 0.968 (0.839 - 1.000) |

### Output of MODEL 2

Haulback mortality rate was higher for large-C than for tuna hook in bigeye tuna, but did not differ among hook types in other species (Table 7, Fig. 4). However, the mortality rate varied with the bait type among species. In bigeye tuna and blue shark, the rate decreased when using fish bait, while in shortfin mako shark, it increased.

**Figure 4.**
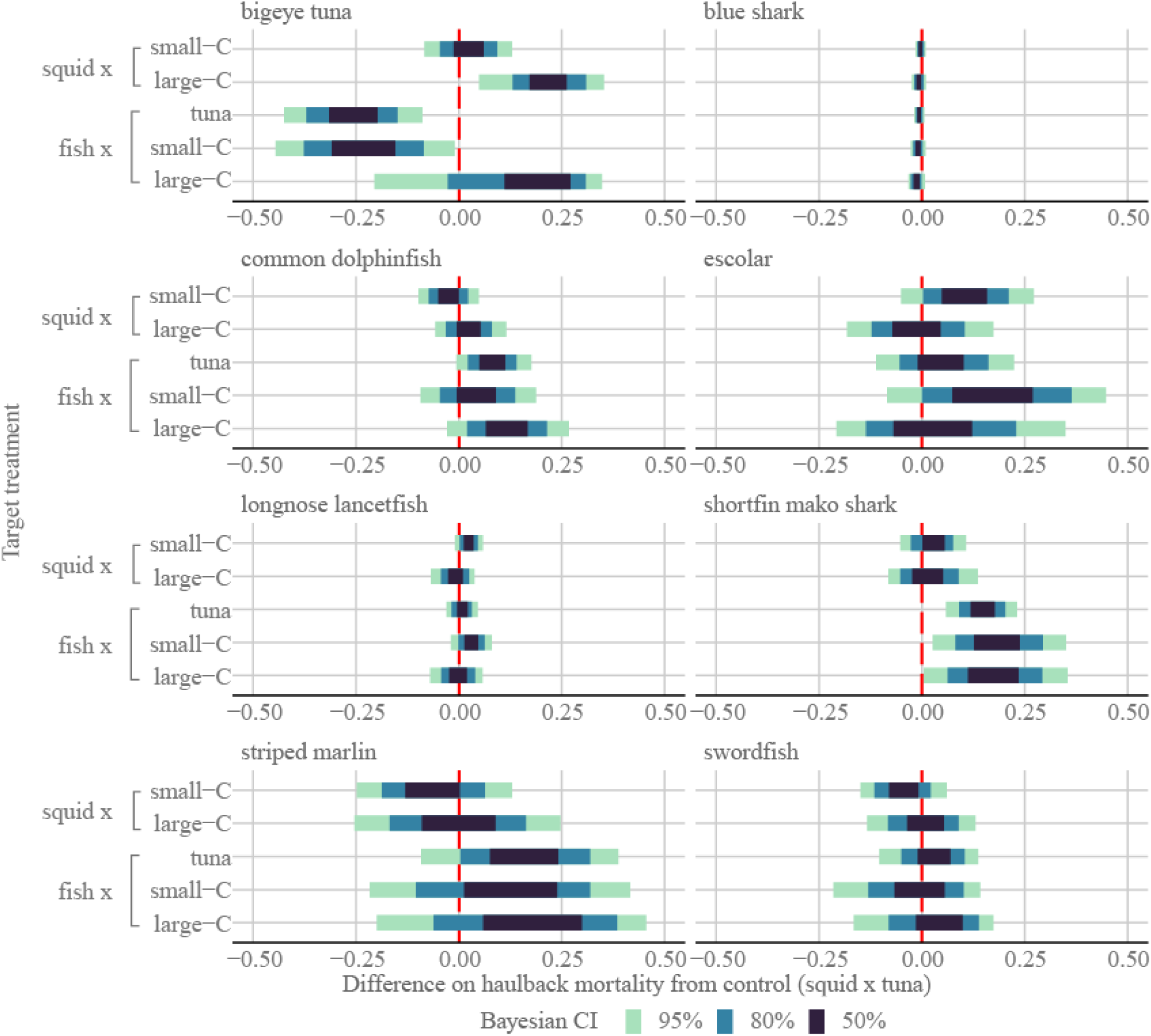
Differences in estimated haulback mortality between each experimental group and the control group (squid x tuna hook). The red dotted line indicates that the difference is zero.

**Table 7.** Haulback mortality rate by hook and bait type (median of posterior distribution). Lower and upper limits of Bayesian credible interval (95% highest density interval [HDI]) are shown in parentheses.

| Species | Hook type |  |  |
| --- | --- | --- | --- |
|  | tuna | small-C | large-C |
| **<u>Bait type: squid</u>** |  |  |  |
| bigeye tuna | 0.734 (0.636 - 0.823) | 0.757 (0.679 - 0.830) | 0.959 (0.813 - 1.000) |
| blue shark | 0.084 (0.077 - 0.093) | 0.081 (0.072 - 0.091) | 0.077 (0.062 - 0.093) |
| common dolphinfish | 0.154 (0.099 - 0.215) | 0.128 (0.081 - 0.180) | 0.180 (0.110 - 0.265) |
| escolar | 0.188 (0.082 - 0.309) | 0.301 (0.165 - 0.454) | 0.182 (0.060 - 0.340) |
| longnose lancetfish | 0.901 (0.870 - 0.930) | 0.925 (0.892 - 0.954) | 0.889 (0.826 - 0.940) |
| shortfin mako | 0.161 (0.106 - 0.223) | 0.187 (0.123 - 0.258) | 0.181 (0.085 - 0.300) |
| striped marlin | 0.532 (0.400 - 0.665) | 0.472 (0.334 - 0.609) | 0.530 (0.318 - 0.740) |
| swordfish | 0.829 (0.754 - 0.895) | 0.783 (0.698 - 0.859) | 0.832 (0.711 - 0.934) |
| **<u>Bait type: fish</u>** |  |  |  |
| bigeye tuna | 0.473 (0.319 - 0.622) | 0.503 (0.337 - 0.663) | 0.884 (0.573 - 1.000) |
| blue shark | 0.078 (0.069 - 0.088) | 0.075 (0.063 - 0.088) | 0.071 (0.056 - 0.088) |
| common dolphinfish | 0.238 (0.159 - 0.326) | 0.201 (0.102 - 0.311) | 0.274 (0.162 - 0.400) |
| escolar | 0.242 (0.114 - 0.390) | 0.371 (0.179 - 0.596) | 0.233 (0.054 - 0.469) |
| longnose lancetfish | 0.909 (0.879 - 0.936) | 0.931 (0.894 - 0.961) | 0.898 (0.841 - 0.947) |
| shortfin mako | 0.307 (0.237 - 0.381) | 0.348 (0.226 - 0.478) | 0.339 (0.196 - 0.493) |
| striped marlin | 0.691 (0.457 - 0.885) | 0.638 (0.375 - 0.870) | 0.691 (0.386 - 0.925) |
| swordfish | 0.853 (0.727 - 0.950) | 0.811 (0.648 - 0.934) | 0.856 (0.700 - 0.967) |

In Fig. 5 and 6 we show the effects of two covariates—sea surface temperature (SST) and soak time—on haulback mortality rate. The response to SST differed by species, with haulback mortality rate increasing with higher SST for common dolphinfish, escolar, shortfin mako sharks and swordfish, and conversely increasing with lower SST for bigeye tuna and longnose lancetfish. We observed little fluctuation in haulback mortality rate with SST in blue shark and striped marlin. In general, haulback mortality rate increased with increasing soak time. However, for bigeye tuna, haulback mortality rate decreased with soak time, but the trend was not clear, and for blue sharks, haulback mortality rate increased only slightly with increased soak time.

**Figure 5.**
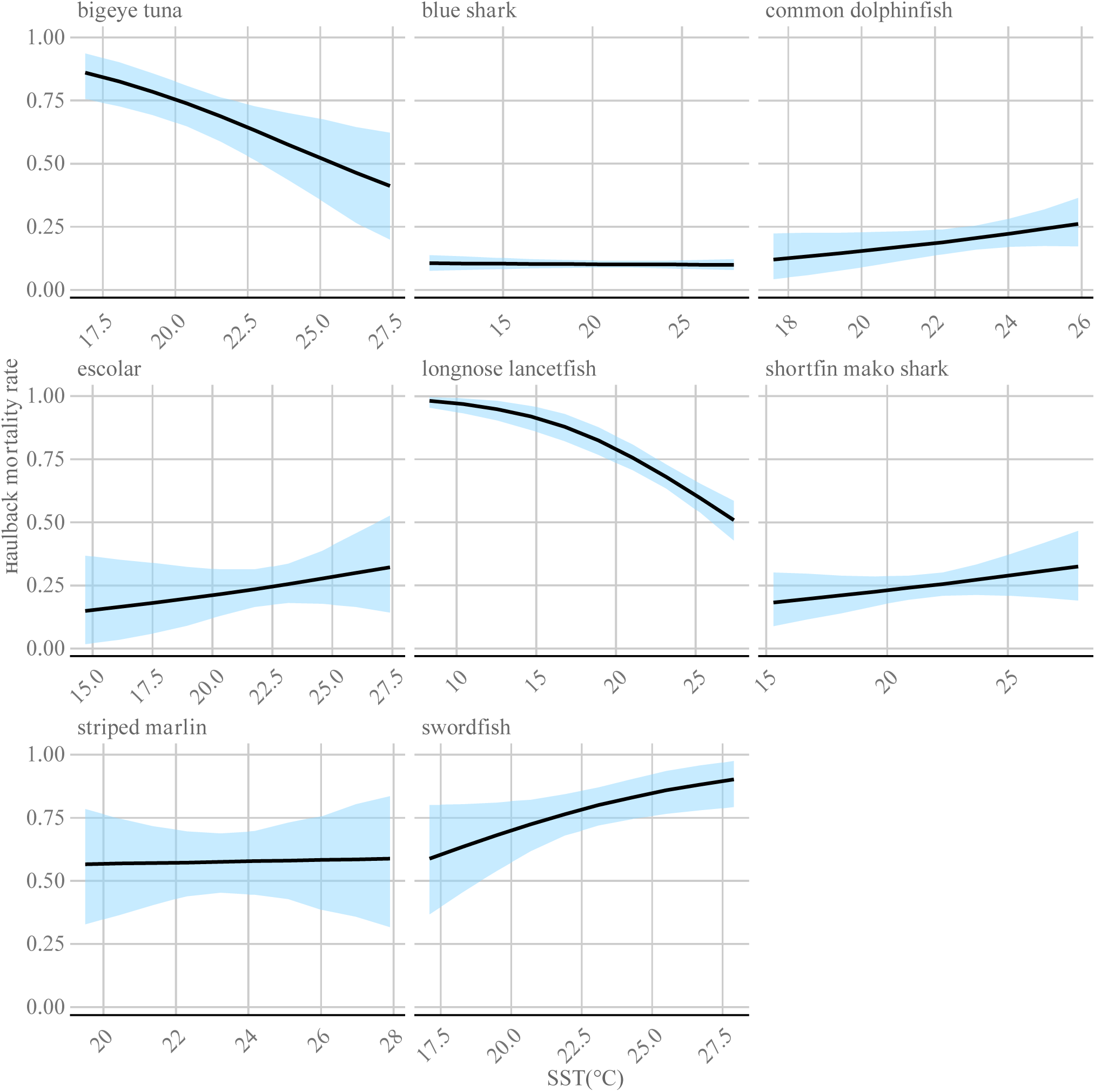
Relationship between sea surface temperature (SST) variability and haulback mortality rate at longline operations. Solid lines indicate median; masked areas indicate 95% Bayesian credible interval.

**Figure 6.**
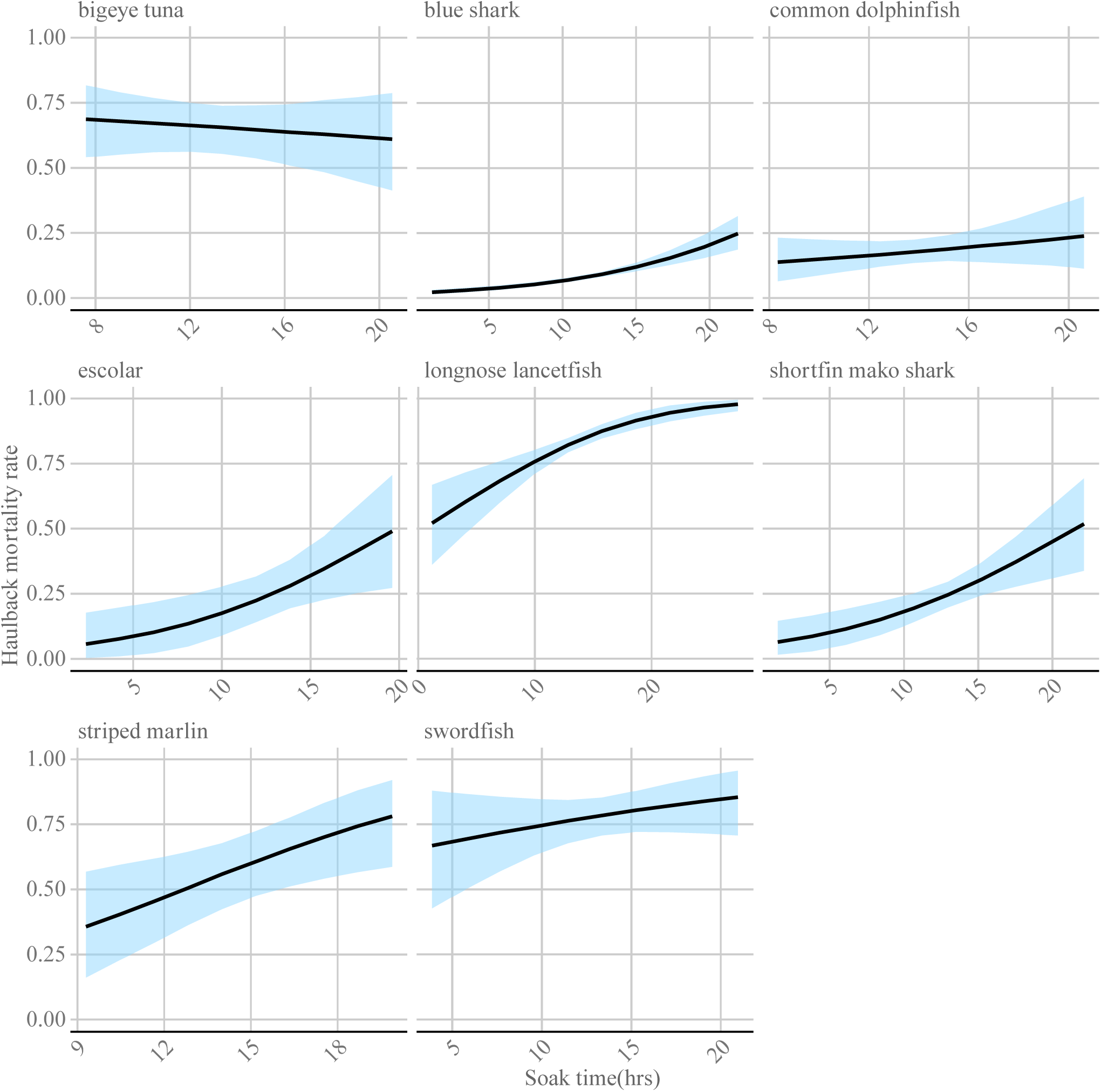
Relationship between soak time (time from setting the branch line to hauling) variability and haulback mortality rate at longline operations. Solid lines indicate median; masked areas indicate 95% Bayesian credible interval.

We observed higher CPUE for only small-C in blue shark, bigeye tuna, common dolphinfish and escolar (Fig. 7, Table 8). We observed differences in standardized CPUE by bait type in bigeye tuna, blue shark, common dolphinfish, escolar, and shortfin mako shark. CPUE decreased with whole fish bait in bigeye tuna and blue sharks, but increased in common dolphinfish, escolar, and shortfin mako shark. In the case of blue shark, escolar, and shortfin mako shark, the bait effect could be varied with circle hooks, suppressing the CPUE-increasing effect by fish bait in bigeye tuna, and blue shark with the use of circle hooks, while conversely this combination boosted increase of CPUE in common dolphinfish, escolar, and shortfin mako shark.

**Figure 7.**
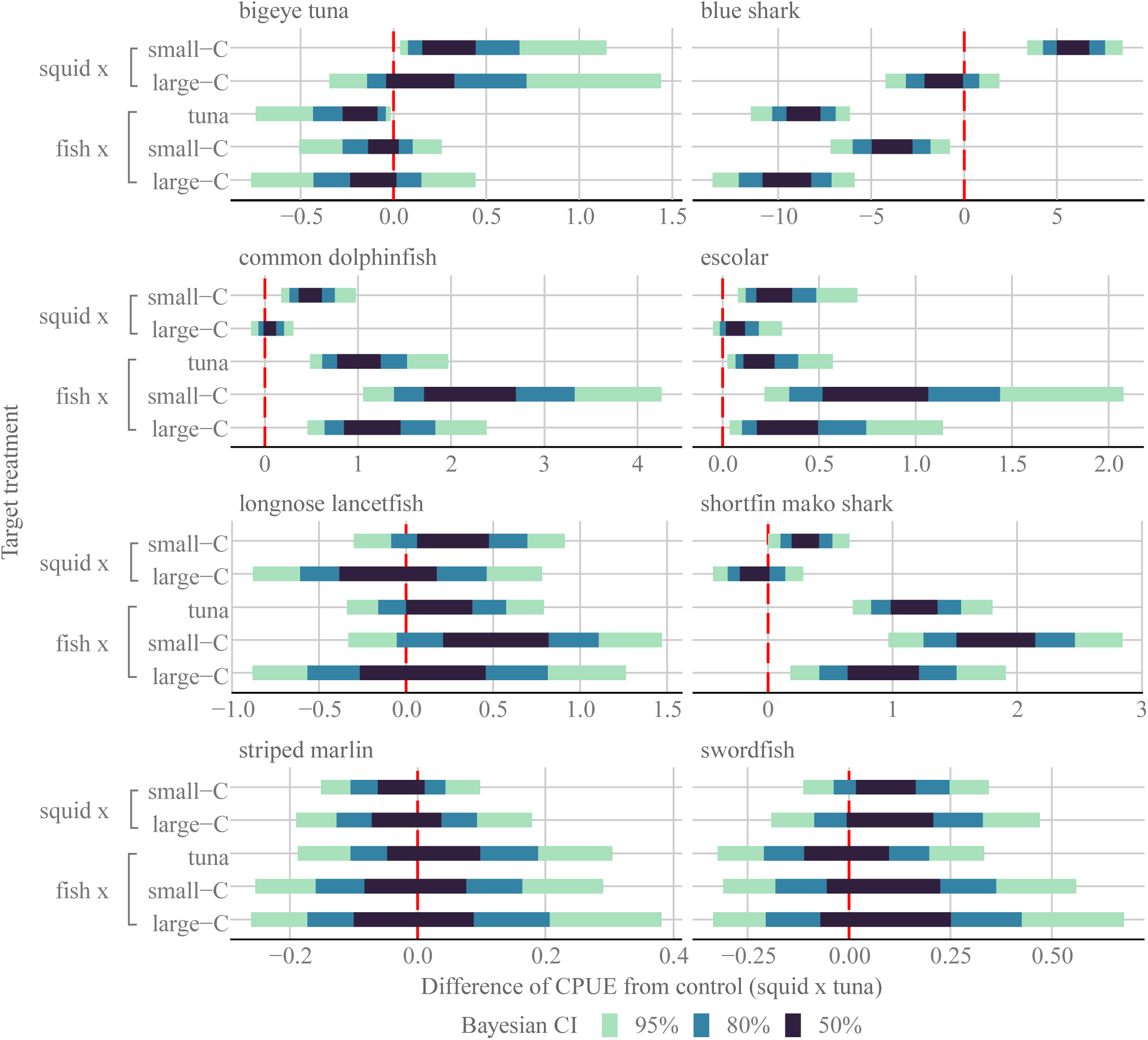
Differences in the standardized catch per unit effort (CPUE) for each experimental group from those for the control group (squid x tuna hook). The red dotted line indicates that the difference is zero.

**Table 8.**
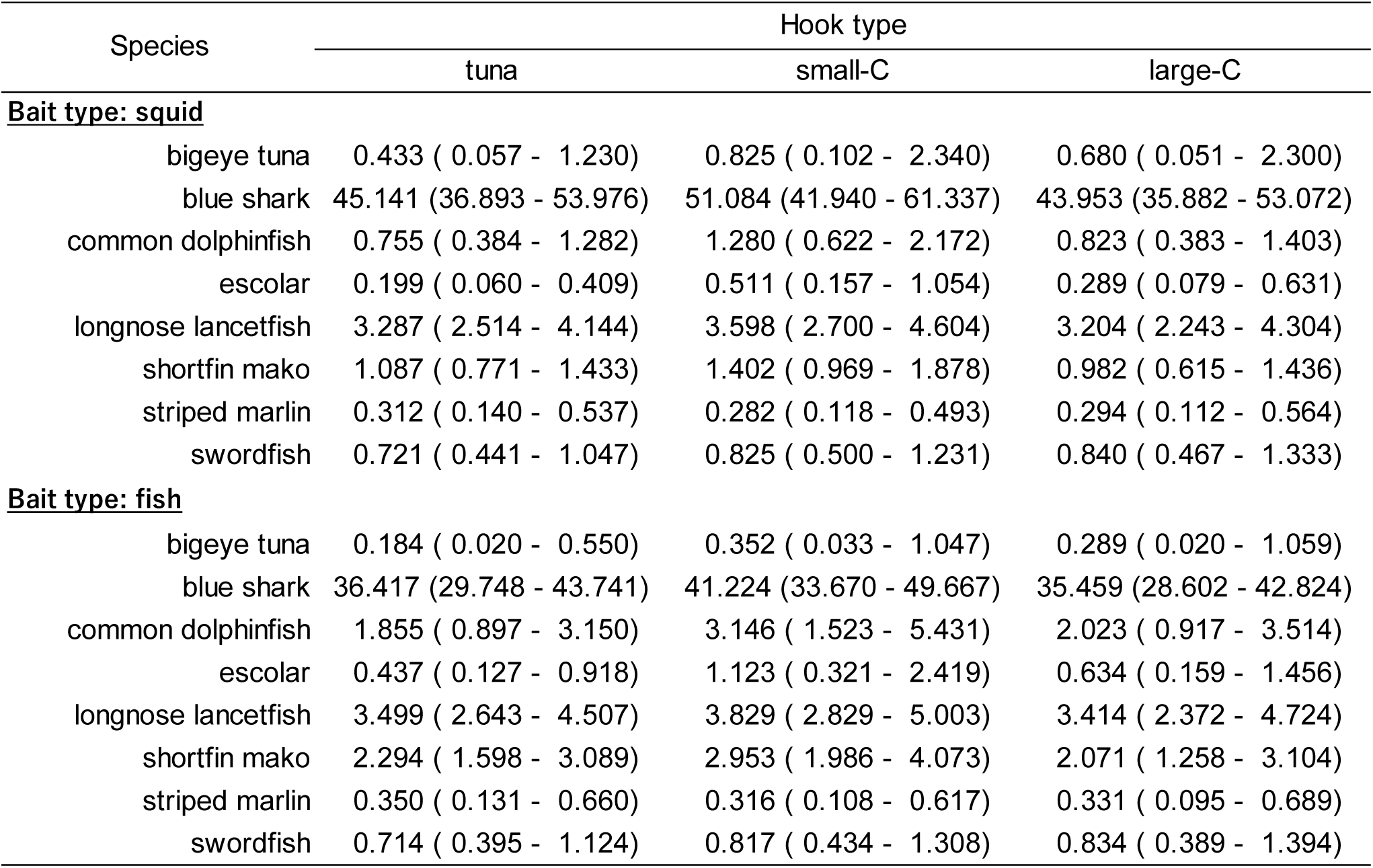
Standardized catch per unit effort (CPUE) by hook and bait type (median of posterior distribution). Lower and upper limits of Bayesian credible interval (95% highest density interval [HDI]) are shown in parentheses.

| Species | Hook type |  |  |
| --- | --- | --- | --- |
|  | tuna | small-C | large-C |
| **<u>Bait type: squid</u>** |  |  |  |
| bigeye tuna | 0.433 ( 0.057 - 1.230) | 0.825 ( 0.102 - 2.340) | 0.680 ( 0.051 - 2.300) |
| blue shark | 45.141 (36.893 - 53.976) | 51.084 (41.940 - 61.337) | 43.953 (35.882 - 53.072) |
| common dolphinfish | 0.755 ( 0.384 - 1.282) | 1.280 ( 0.622 - 2.172) | 0.823 ( 0.383 - 1.403) |
| escolar | 0.199 ( 0.060 - 0.409) | 0.511 ( 0.157 - 1.054) | 0.289 ( 0.079 - 0.631) |
| longnose lancetfish | 3.287 ( 2.514 - 4.144) | 3.598 ( 2.700 - 4.604) | 3.204 ( 2.243 - 4.304) |
| shortfin mako | 1.087 ( 0.771 - 1.433) | 1.402 ( 0.969 - 1.878) | 0.982 ( 0.615 - 1.436) |
| striped marlin | 0.312 ( 0.140 - 0.537) | 0.282 ( 0.118 - 0.493) | 0.294 ( 0.112 - 0.564) |
| swordfish | 0.721 ( 0.441 - 1.047) | 0.825 ( 0.500 - 1.231) | 0.840 ( 0.467 - 1.333) |
| **<u>Bait type: fish</u>** |  |  |  |
| bigeye tuna | 0.184 ( 0.020 - 0.550) | 0.352 ( 0.033 - 1.047) | 0.289 ( 0.020 - 1.059) |
| blue shark | 36.417 (29.748 - 43.741) | 41.224 (33.670 - 49.667) | 35.459 (28.602 - 42.824) |
| common dolphinfish | 1.855 ( 0.897 - 3.150) | 3.146 ( 1.523 - 5.431) | 2.023 ( 0.917 - 3.514) |
| escolar | 0.437 ( 0.127 - 0.918) | 1.123 ( 0.321 - 2.419) | 0.634 ( 0.159 - 1.456) |
| longnose lancetfish | 3.499 ( 2.643 - 4.507) | 3.829 ( 2.829 - 5.003) | 3.414 ( 2.372 - 4.724) |
| shortfin mako | 2.294 ( 1.598 - 3.089) | 2.953 ( 1.986 - 4.073) | 2.071 ( 1.258 - 3.104) |
| striped marlin | 0.350 ( 0.131 - 0.660) | 0.316 ( 0.108 - 0.617) | 0.331 ( 0.095 - 0.689) |
| swordfish | 0.714 ( 0.395 - 1.124) | 0.817 ( 0.434 - 1.308) | 0.834 ( 0.389 - 1.394) |

Compared to differences in CPUE among hook and bait type, those in MPUE were relatively small (Table 9). We observed differences with higher MPUE for only small-C in bigeye tuna and escolar, (Fig. 8). We confirmed decreases in MPUE by fish bait in bigeye tuna and blue shark, and conversely, increases in MPUE in common dolphinfish, shortfin mako shark. The effect of the combination of whole fish bait and circle hook varied in bigeye tuna, blue shark, common dolphinfish, escolar and shortfin mako shark. In bigeye tuna and blue shark, the effect of the circle hook on MPUE was suppressed by fish bait, while in common dolphinfish, escolar and shortfin mako shark, MPUE increased significantly when whole fish bait and circle hook were used together.

**Figure 8.**
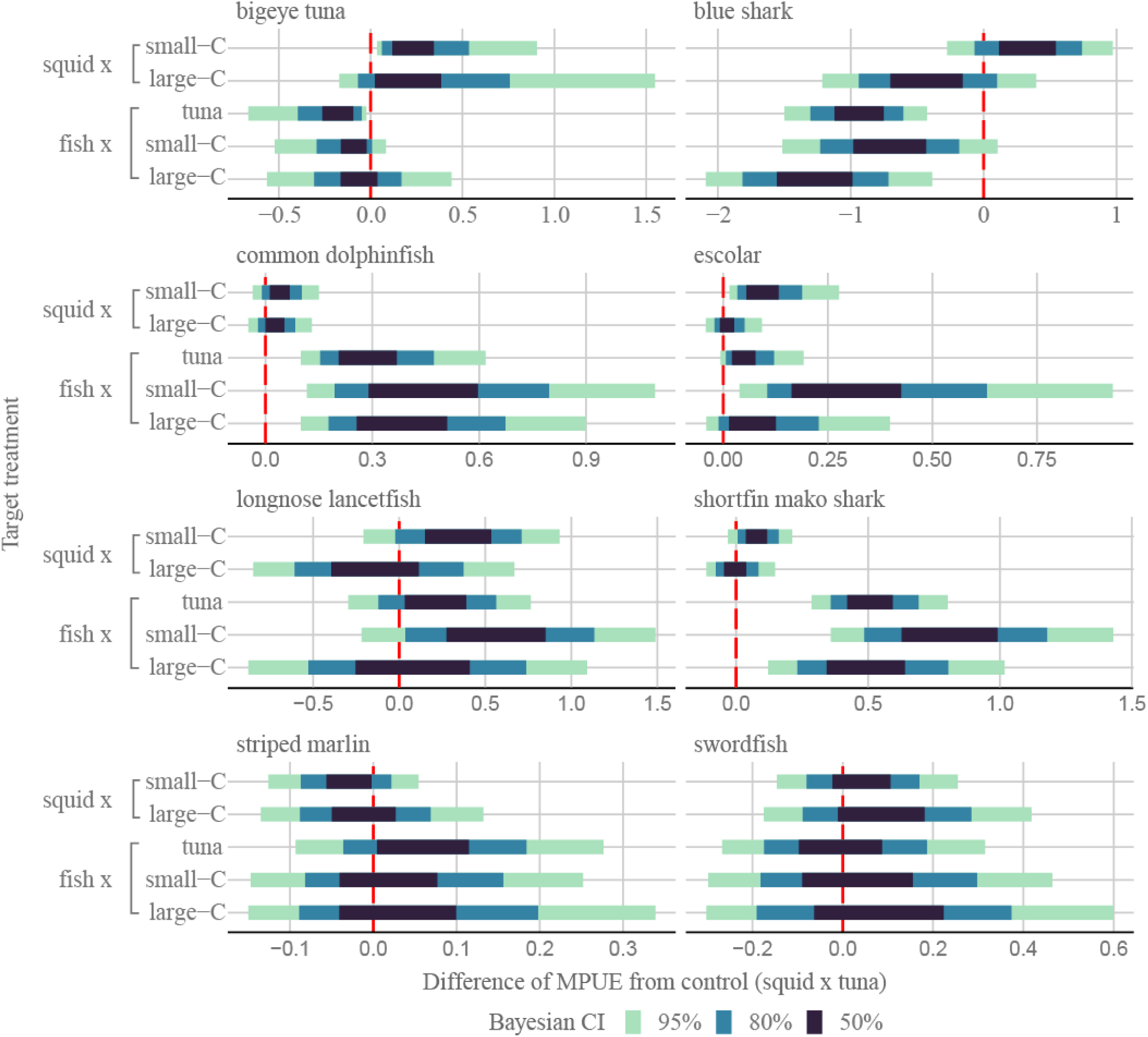
Differences in the estimated MPUE (mortality per unit effort) between those for each experimental group and those for the control group (squid x tuna hook). The red dotted line indicates that the difference is zero.

**Table 9.** Estimated MPUE by hook and bait type (median of posterior distribution). Lower and upper limits of Bayesian credible interval (95% highest density interval [HDI]) are shown in parentheses.

| Species | Hook type |  |  |
| --- | --- | --- | --- |
|  | tuna | small-C | large-C |
| **<u>Bait type: squid</u>** |  |  |  |
| bigeye tuna | 0.316 (0.035 - 0.899) | 0.623 (0.078 - 1.774) | 0.637 (0.051 - 2.173) |
| blue shark | 3.802 (3.031 - 4.628) | 4.143 (3.263 - 5.120) | 3.383 (2.494 - 4.401) |
| common dolphinfish | 0.115 (0.048 - 0.211) | 0.163 (0.067 - 0.301) | 0.148 (0.056 - 0.279) |
| escolar | 0.037 (0.008 - 0.088) | 0.152 (0.035 - 0.345) | 0.052 (0.007 - 0.140) |
| longnose lancetfish | 2.960 (2.255 - 3.734) | 3.322 (2.457 - 4.225) | 2.839 (1.986 - 3.849) |
| shortfin mako | 0.174 (0.096 - 0.261) | 0.260 (0.144 - 0.397) | 0.177 (0.066 - 0.328) |
| striped marlin | 0.165 (0.068 - 0.294) | 0.132 (0.051 - 0.243) | 0.153 (0.048 - 0.320) |
| swordfish | 0.595 (0.353 - 0.866) | 0.643 (0.388 - 0.971) | 0.694 (0.369 - 1.109) |
| **<u>Bait type: fish</u>** |  |  |  |
| bigeye tuna | 0.087 (0.008 - 0.266) | 0.174 (0.017 - 0.540) | 0.241 (0.014 - 0.906) |
| blue shark | 2.844 (2.232 - 3.519) | 3.094 (2.359 - 3.935) | 2.527 (1.816 - 3.348) |
| common dolphinfish | 0.438 (0.176 - 0.800) | 0.627 (0.213 - 1.238) | 0.550 (0.197 - 1.057) |
| escolar | 0.104 (0.019 - 0.250) | 0.407 (0.086 - 0.996) | 0.144 (0.015 - 0.439) |
| longnose lancetfish | 3.177 (2.389 - 4.094) | 3.558 (2.611 - 4.651) | 3.055 (2.075 - 4.207) |
| shortfin mako | 0.703 (0.444 - 1.004) | 1.020 (0.547 - 1.586) | 0.697 (0.312 - 1.196) |
| striped marlin | 0.237 (0.075 - 0.469) | 0.195 (0.054 - 0.413) | 0.219 (0.051 - 0.497) |
| swordfish | 0.601 (0.323 - 0.960) | 0.653 (0.324 - 1.065) | 0.703 (0.320 - 1.201) |

### Hook and bait effect for sea turtle bycatch

We failed to complete our analysis for loggerhead turtles throughout the models because the bias in frequency of capture events among the experimental groups was too large and did not converge except for only a part of MODEL 1. Instead, we show the nominal CPUE, haulback mortality rate, and MPUE for each experimental group in Table 10, and hooking location and haulback mortality rate by each hook type with squid bait in Fig. 9. Most individuals survived regardless of hooking location. When whole fish bait was used, haulback mortality rate and associated MPUE were zero. For squid bait, large-C had the smallest CPUE, mortality, and MPUE.

**Figure 9.**
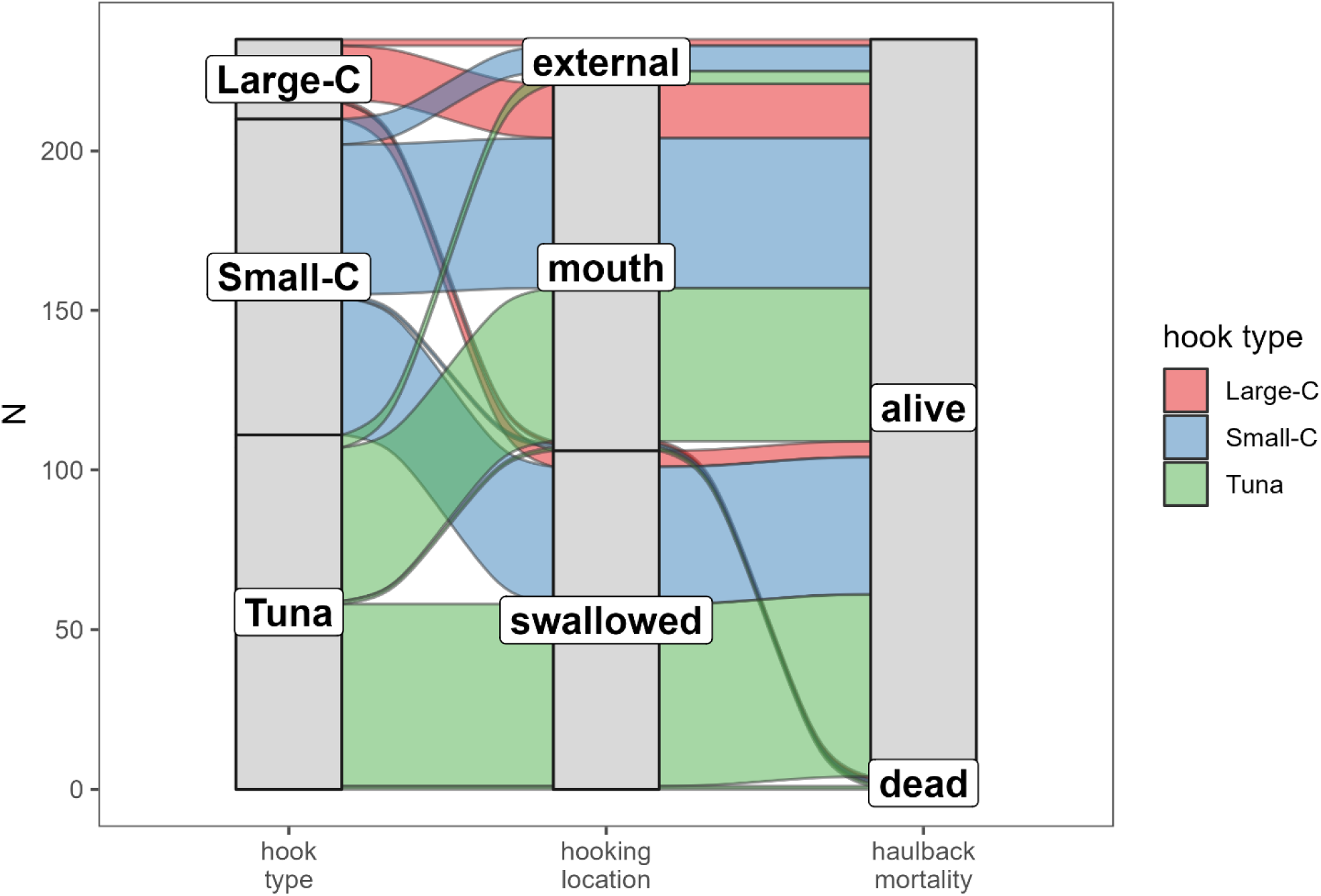
Alluvial plot of hooking locations and associated haulback mortality rates of loggerhead turtles by hook when squid bait is used.

**Table 10.**
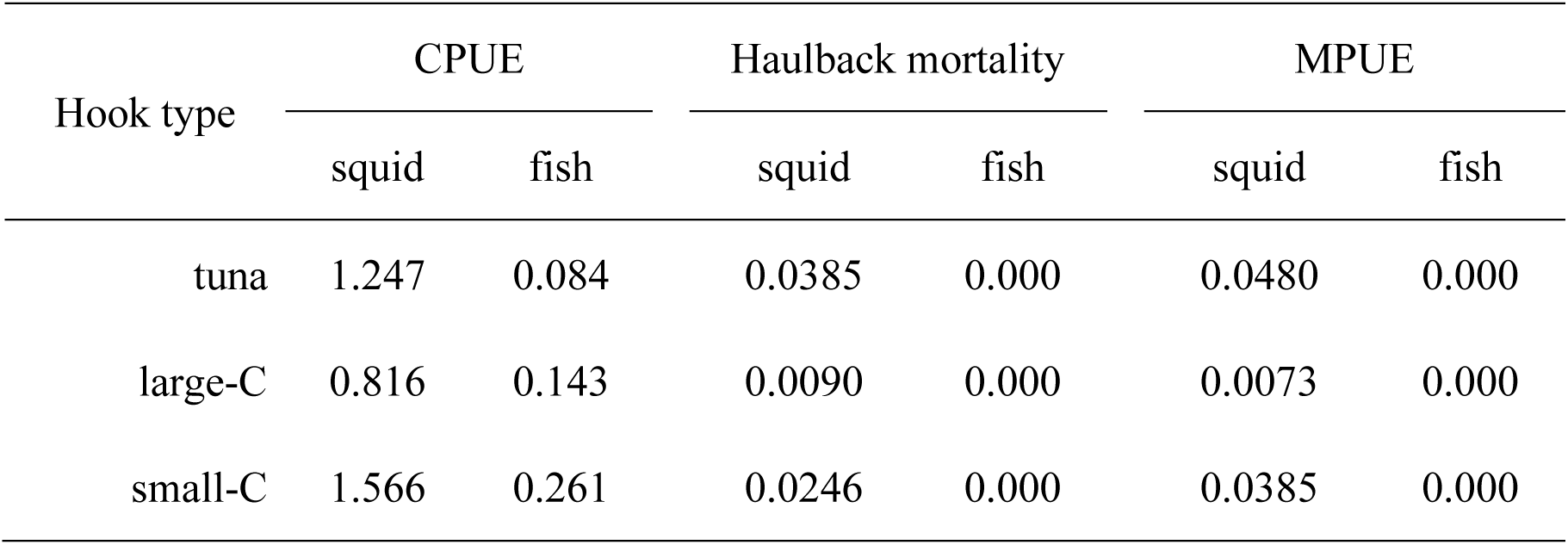
Nominal CPUE, Haulback mortality rate and MPUE of loggerhead turtle and all figures are based on aggregated operational data, not estimates.

## DISCUSSION

Our experimental comparisons showed that the hook and bait type—both considered as effective bycatch mitigation measures for sea turtles—have extremely multifaceted effects for teleost fishes and sharks and, in some species, the direction of the effects was conflicted. The results provide significant insight into two aspects of the management of vulnerable bycatch species in tuna fisheries: how the confrontational effect of bycatch mitigation measures should be managed, and in which processes of fishing mortality intervention in the management of vulnerable species should occur. As a specific concern regarding the former, when considering shortfin mako shark, which are experiencing significant stock depletion in the North Atlantic (Sims et al. 2018; ICCAT 2019), the implementation of bycatch mitigation measures for sea turtles, which are also required to reduce fishery bycatch, could conversely increase fishing mortality and become a conservation risk. Previously, most of bycatch mitigation measure assessments have focused on whether they reduce the impact of vulnerable bycatch species of concern being bycaught, with the secondary impact being from an economic perspective—in other words, whether the catch rate of commercial species is reduced or not. Where both loggerhead turtle and shortfin mako with opposing effects are at low abundance, management measures should be based on a thorough discussion identifying the optimal combination of mitigation measures, accompanied by scientific evidence. With regard to the latter issue, the mortality reduction expected from circle hooks is not very promising, especially for species with high catch mortality, since the main effect of circle hooks is to minimize internal organ damage, which is of little use for species that have died from other causes, such as heat stress or suffocation. For such species, consideration of methods to reduce the catch itself rather than the mortality rate will be of greater benefit.

Increased mouth hooking of loggerhead turtle by large circle hooks is consistent with existing studies. The circle hook prevents internal organ damage and improves the probability of live release (Cooke & Suski 2004) while the impact of circle hooks on haulback mortality in this study could not be evaluated due to skewed data about mortality events. For the same reason, the MODEL 2 analysis could not evaluate the effects of circle hook and bait type on CPUE, mortality rate, and MPUE of loggerhead turtle. However, since mortality event of loggerhead turtle did not occur at all when fish bait was used, it may be assumed that there is an effect of mortality reduction by using fish bait. This result is also consistent with existing studies, and is related to the lower attractant effect of whole fish bait on marine turtles and the increased probability of swallowing caused by the difficulty to bite off the bait (Stokes et al. 2011; Parga et al. 2015).

Our study indicated that reduced hook swallowing with circle hooks and increased haulback mortality after hook swallowing were found for sharks and swordfish. Hook swallowing has been reported to increase the likelihood of fatal damage to internal organs. Although previous studies have reported that studies to attach satellite tags to white marlin *Kajikia albida* caught by recreational fishing using circle hooks and subsequently released have reduced the post-release mortality rate (Horodysky & Graves 2005), unfortunately, the present study did not corroborate this information. Our results also presented different effects of hooking location by hook types, with more frequent mouth hooking by small circle hooks in many species. Few studies have examined the relationship among size of circle hook, hooking location and haulback mortality of non-turtle species. However, two studies discussed the possibility that relative differences in mouth and hook size, and differences in feeding behavior toward prey (swallowing the prey whole or biting it off) may affect the hooking location (Epperly et al. 2012; Gilman et al. 2020).

An increase in CPUE and MPUE was observed only with small circle hooks among the two types of circle hooks. This indicates that the increase in CPUE due to small circle hooks had a greater impact on MPUE than the haulback mortality rate. There have been many previous findings on the effects of circle hook use on CPUE, with elevated CPUE for tunas and no consistent trend for other teleosts and sharks. Interestingly, we did not observe an increased CPUE and MPUE with large circle hooks. Although few previous studies have focused on hook size and made comparisons, catch rates for skipjack, shortbill spearfish, escolar, and lancetfish are reported to have decreased when larger hooks were used (Curran & Beverly 2012; Gilman et al. 2018). Considering the effect of hook size in terms of the catch process, it is unlikely that catch rates increased due to swallowing, as the results of MODEL 1 indicate an increase in mouth hooking for many species. In the case of blue shark and shortfin mako shark, the increase was observed in CPUE for small circle hook, but this increase was not observed in MPUE. It may be explained by that the effect of the circle hook on MPUE may have been masked by the uncertainty of haulback mortality rate. Several studies have examined the effects of fish bait without circle hook and have reported reduced catch rates for tropical tunas, blue sharks, and escolar and increased catch rates for shortfin mako, porbeagle shark, and white marlin (Fernandez-Carvalho et al. 2015; Foster et al. 2012; Watson et al. 2005; Yokota et al. 2009). In swordfish, some previous studies evaluating the effect of switching to whole fish bait from squid bait have reported conflicting effects (increase: Santos et al. 2012; Foster et al. 2012; decrease: Fernandez-Carvalho et al. 2015) but few studies have referred to haulback mortality rate by bait types other than those on sea turtles. The catch rates of the target species, as previously noted, were affected by bait texture, but a mechanistic explanation for mortality effects is lacking. Since the likelihood that differences in feeding behavior among species have an effect is high, this issue could be resolved through comparative studies based on observations of feeding behavior, as in the case of circle hooks.

When the effects of hook and bait types were considered simultaneously, it was found that the bait type had a more significant impact on CPUE, mortality rate, and MPUE than the hook type. However, whether this impact was beneficial or detrimental varied greatly depending on the species. Although the combination of circle hooks and fish bait is effective for avoiding sea turtle bycatch, this combination may pose a high mortality risk for endangered species like shortfin mako shark, and even in the case of target fishes like bigeye tuna, may counteract the expected positive effect of the circle hook on catch rate. In the case of shortfin mako, for example, changing the bait from squid to fish increases MPUE by about 4.0 times, and changing from tuna hooks to small circle hooks further increases MPUE by about 5.9 times (Table 9), and in the case of bigeye tuna, the CPUE estimate, which increased by 1.9 times with small circle hooks, returned to the same level as tuna hooks by changing from squid to fish bait (Table 8). Such substantial changes in CPUE and MPUE would not be ignored when managing fisheries for those species. Although very limited studies have simultaneously examined the interrelationship between hook and bait types, all studies support the conclusion that the combination of hook type and bait type causes fluctuations in catch rates and that the direction of response varies among species (Coelho et al. 2012; Foster et al. 2012; Fernandez-Carvalho et al. 2015).

Water temperature and soak time emerged as significant factors affecting haulback mortality rate, which had been reported in sharks (Carruthers et al. 2009; Gallagher et al. 2014) and sea turtles (Watson et al. 2005). This indicates these covariates need to be controlled statistically or experimentally when assessing the effects of hook and bait type on mortality rate. The effect of water temperature—particularly during the depth and time of day when hooked—and changes in water temperature up to the time the fish is landed, are considered to be influential. In addition, in high water temperature environments, studies have identified an increased risk of suffocation due to decreased dissolved oxygen in water and increased physiological metabolic rate (Skomal & Bernal 2010). Gallagher et al. (2014) reported an increase in haulback mortality rate for four shark species when caught during high water temperatures. In addition, for the species that adopt rum ventilation, prolonged soak time inevitably increases the risk of suffocation due to the restriction of swimming behavior by being hooked. Mortality rates of tuna, swordfish, and sharks reportedly increased with increasing soak time (Epperly et al. 2012; Gallagher et al. 2014).

We quantified our data through experimental operations that standardized the various conditions, but not all aspects were completely controlled. For example, while previous studies on hook size have examined the correspondence with actual measurements (Gilman et al. 2016), several shapes of circle hook were used in the experiment in this study, precluding examination of effects of individual hook types due to sample size issues. We were also unable to examine hooking location of the catch when fish bait was used. These omissions, while having a limited impact on the present conclusions, are probably variables that should be considered for a deeper examination of the effects of terminal gear on catch and bycatch. In this experiment, wire leaders were used on all branchlines to minimize the effects of sharks’ bite-off. While some studies have described concerns that wire leaders may increase catch rates, especially for rare sharks, they are considered essential for at least experimentally verifying accurate catch and mortality rates for shark species. We know from this and previous studies that haulback mortality rates for sharks are much lower than those for teleosts (Afonso et al. 2012; Reinhardt et al. 2018), and the implementation of safe release protocols, even with wire leaders, allow for the reduction of risk for vulnerable shark species.

Here, based on a Bayesian approach, we succeeded in presenting a quantitative impact assessment of terminal gear on teleosts, sharks, and sea turtles by directly calculating the expected values for mortality rate, CPUE, and MPUE with each terminal gear. Calculating MPUE using this model can be a very useful tool because it provides a more direct estimate than does CPUE or mortality rate alone of catch/bycatch risk to populations of those species. Although we did not include post-release mortality rate in the model due to lack of data, it would be possible to estimate overall fishing mortality in the model by designing additional experiments so that mark–recapture is conducted at the same time. Even if it is not possible to use wire leaders for the proportion of “cryptic catch” due to bite-off, it is possible to extrapolate this proportion into the model to make predictions regarding mortality—a development we anticipate. Although the data used in the analysis relied solely on the results of an Asian-style longline experiment in the Pacific Ocean and may therefore contain inherent biases, the same analysis method can be used in conjunction with data from other experiments conducted in other areas and fishing styles to provide a more integrated assessment.

---

**Appendix S1.**
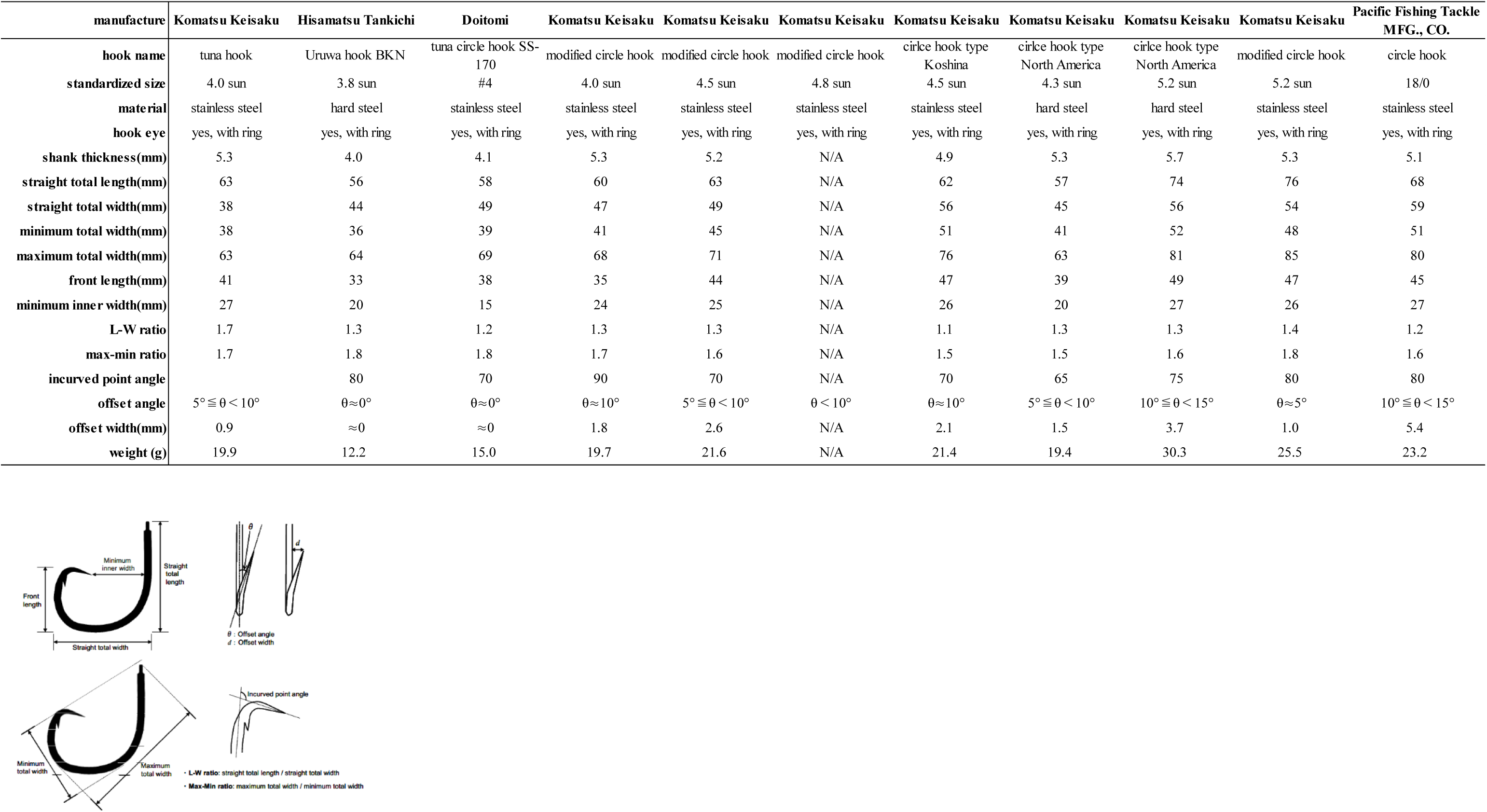
Detailed measurements of hooks used in the experiment and a figure explaining measurement points of hooks (copied from Yokota et al. 2006b).

**Appendix S2.**
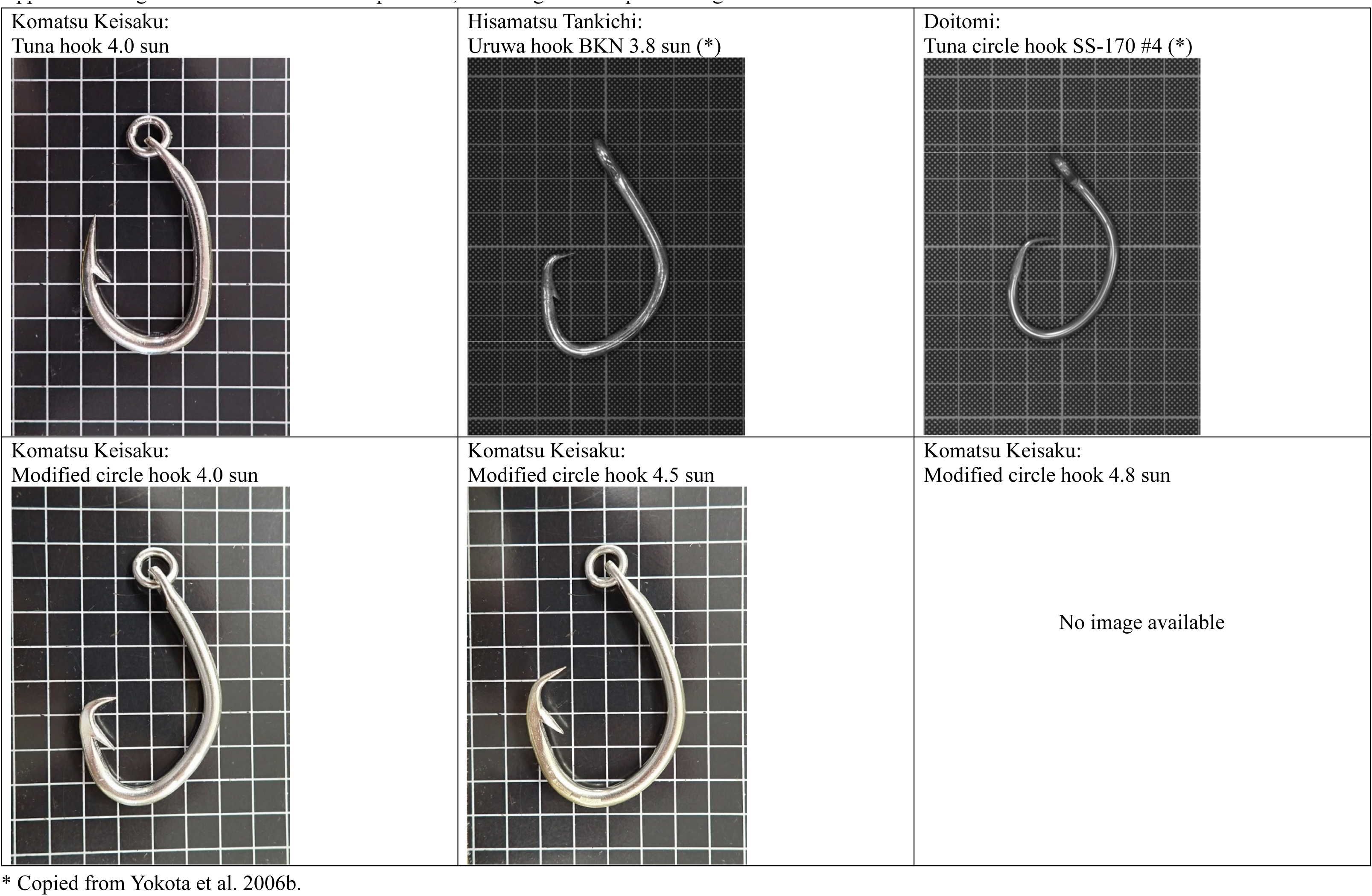
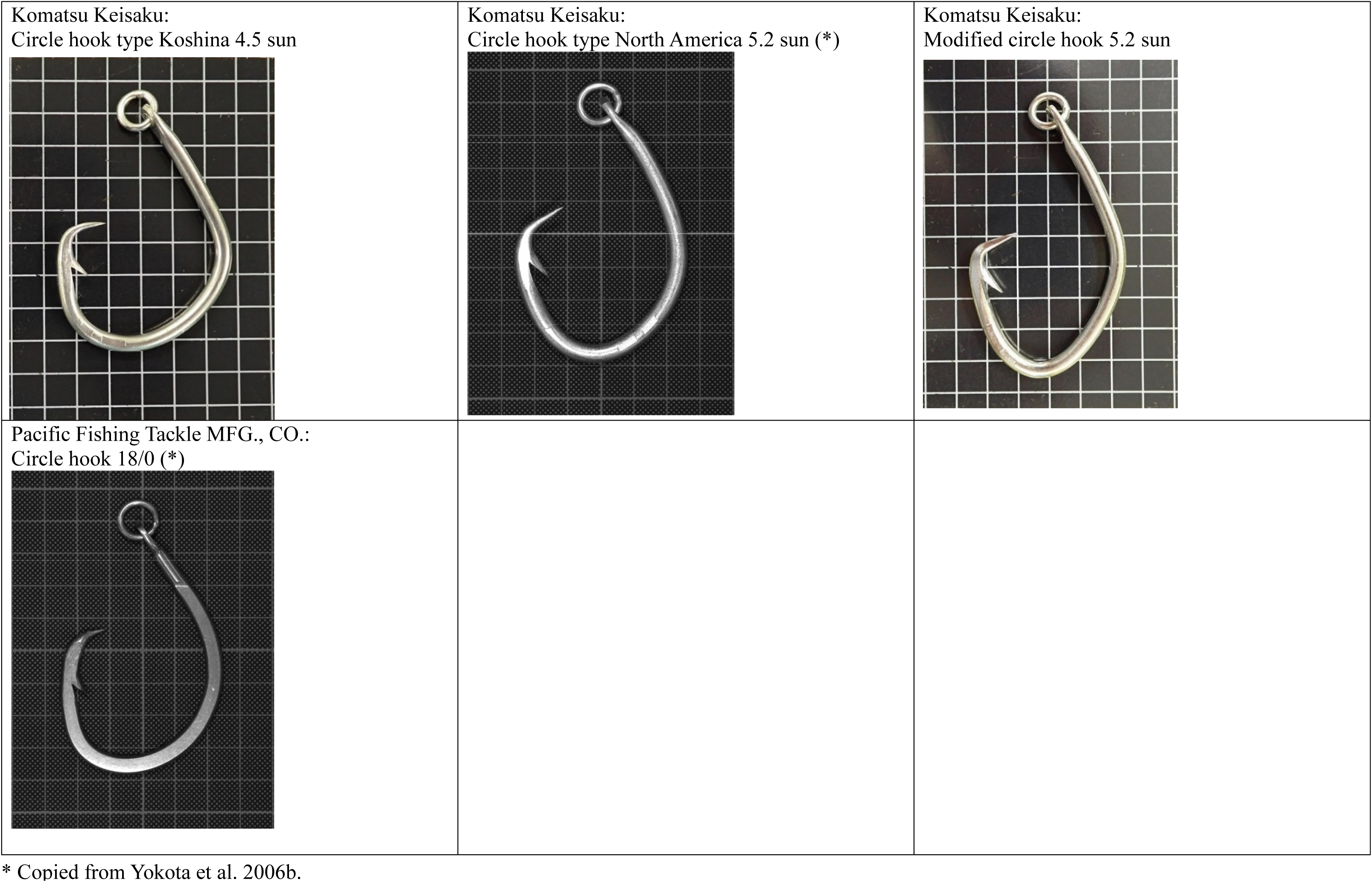
Figure of hooks used in the experiment, including a 1 cm square background.

**Appendix S3.**
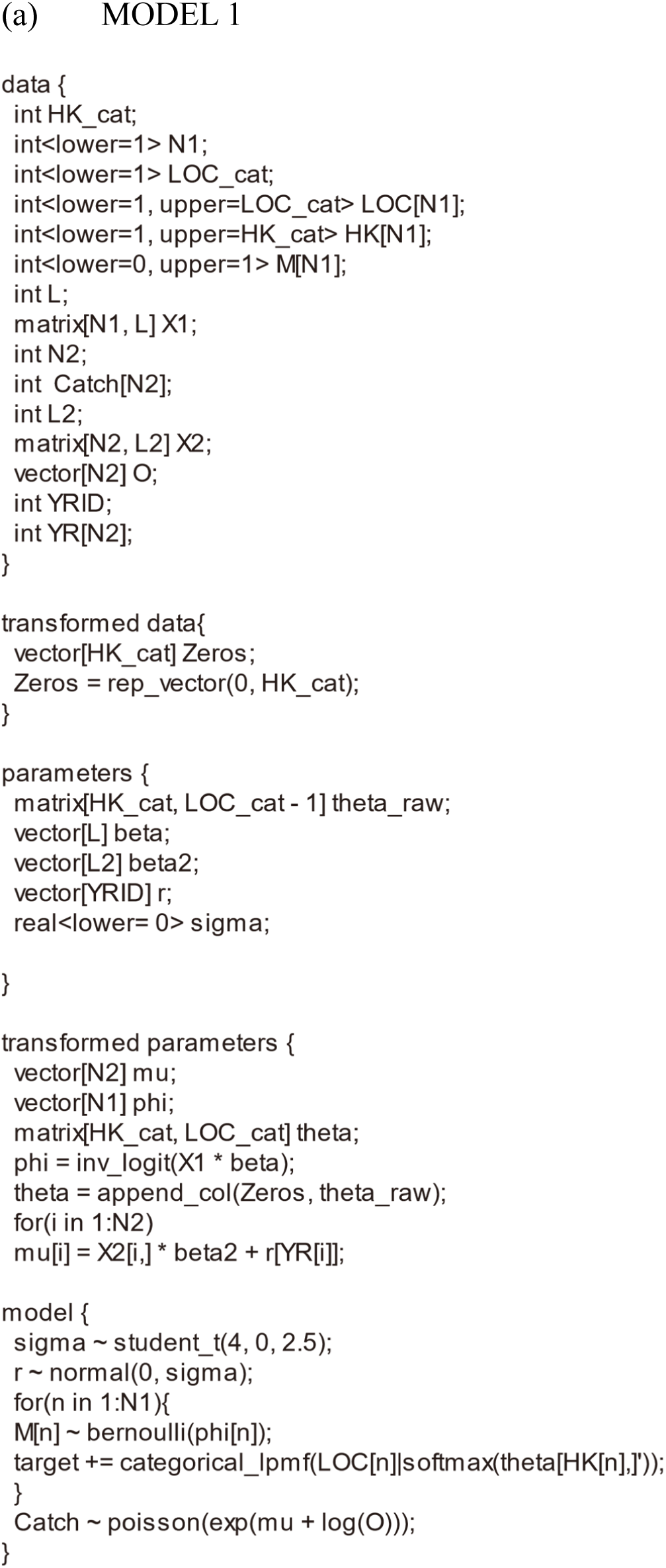
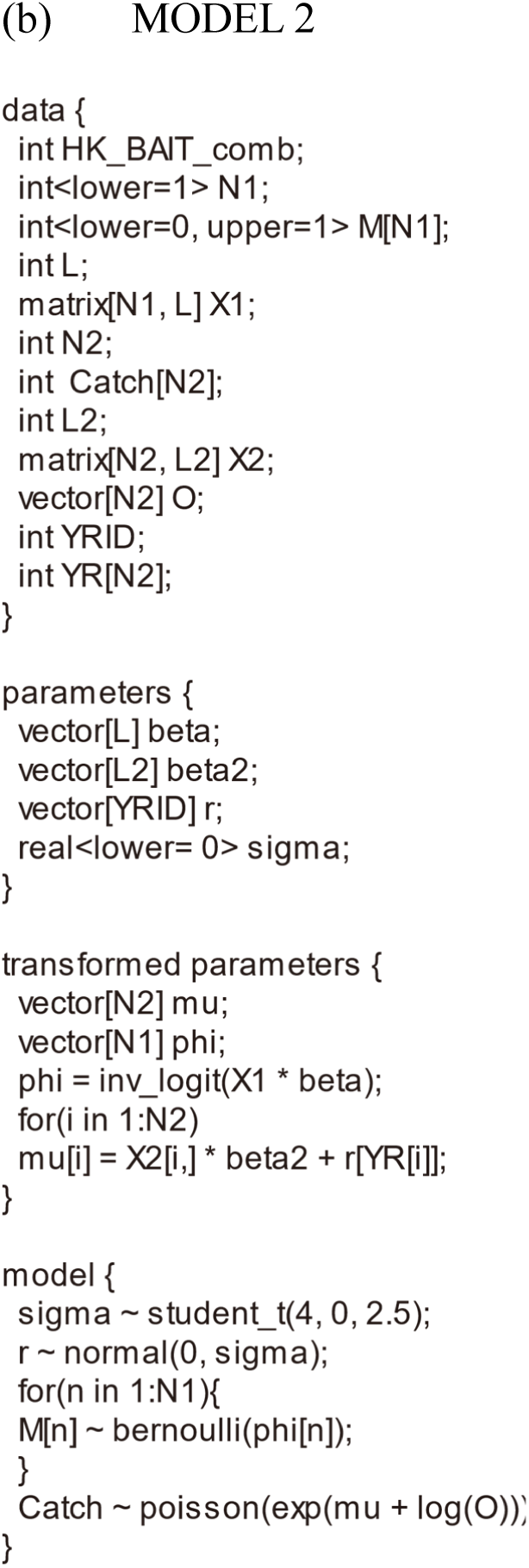
Stan code used for MCMC sampling of (a) Model 1 and (b) Model 2

**Appendix S4.**
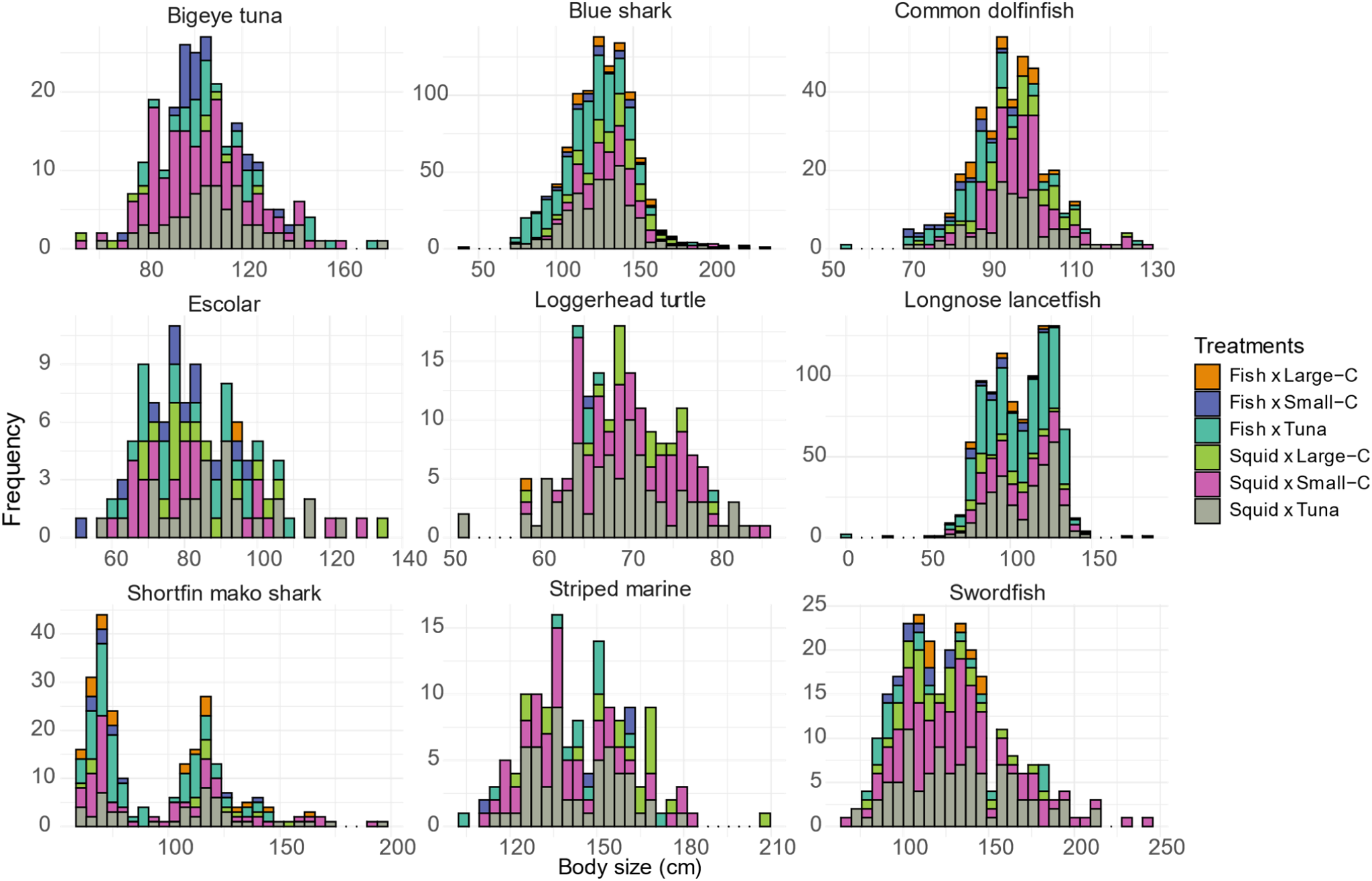
Size distributions by species for the major species captured in the study, with body length as an index of precaudal length for blue sharks and shortfin mako sharks, straight-line carapace length for loggerhead turtles, eye-to-fork length for striped marlin and swordfish, and fork length for all other species.

## Notes

### Competing Interest Statement

The authors have declared no competing interest.

### Summary of Updates

Corrected a contradictory description in the Results section. Updated author information.

## REFERENCES

Afonso AS, Hazin FHV, Carvalho F, Pacheco JC, Hazin H, Kerstetter DW, Murie D, Burgess GH (2011) Fishing gear modifications to reduce elasmobranch mortality in pelagic and bottom longline fisheries off Northeast Brazil. Fisheries Research 108, 336–343.

Afonso AS, Santiago R, Hazin H, Hazin FHV (2012) Shark bycatch and mortality and hook bite-offs in pelagic longlines: Interactions between hook types and leader materials. Fish Res. 131–133, 9–14. 10.1016/j.fishres.2012.07.001.

Andraka S, Mug M, Hall M, Pons M, Pacheco L, Parrales M, Rendón L, Parga ML, Mituhasi T, Segura Á, et al. (2013) Circle hooks: Developing better fishing practices in the artisanal longline fisheries of the Eastern Pacific Ocean. Biological Conservation 160, 214–224. 10.1016/j.biocon.2013.01.019

Arnold JB (2021) ggthemes: Extra Themes, Scales and Geoms for “ggplot2.” Available at https://cran.r-project.org/package=ggthemes

Becker RA, Wilks AR, Brownrigg R (2020) mapdata: Extra Map Databases. Available at https://cran.r-project.org/package=mapdata

Becker RA, Wilks AR, Brownrigg R, Minka TP, Deckmyn A (2022) maps: Draw geographical maps. Available at https://cran.r-project.org/package=maps

Brunson JC (2020) ggalluvial: Layered Grammar for Alluvial Plots. Journal of Open Source Software 5:2017.

Carruthers EH, Schneider DC, Neilson, JD (2009) Estimating the odds of survival and identifying mitigation opportunities for common bycatch in pelagic longline fisheries. Biological Conservation 142, 2620–2630. 10.1016/j.biocon.2009.06.010

Clarke S, Sato M, Small C, Sullivan B, Inoue Y, Ochi D (2015) Bycatch in longline fisheries for tuna and tuna-like species; A global review of status and mitigation measures. FAO Fisheries and Aquaculture Technical Paper 588. Available at https://www.fao.org/3/a-i4017e.pdf.

Coelho R, Santos MN, Amorim S (2012) Effects of hook and bait on targeted and bycatch fishes in an equatorial Atlantic pelagic longline fishery. Bulletin of Marine Science 88, 449–467.

Cooke SJ, Suski CD (2004) Are circle hooks an effective tool for conserving marine and freshwater recreational catch-and-release fisheries? Aquatic Conservation: Marine and Freshwater Ecosystems 14, 299–326.

Curran D, Beverly S (2012) Effects of 16/0 circle hooks on pelagic fish catches in three south pacific albacore longline fisheries. Bulletin of Marine Science 88, 485–497. 10.5343/BMS.2011.1060

Dias MP, Martin R, Pearmain EJ, Burfield IJ, Small C, Phillips RA, Yates O, Lascelles B, Borboroglu PG, Croxall JP (2019) Threats to seabirds: A global assessment. Biological Conservation 237, 525–537.

Diaz GA (2008) The effect of circle hooks and straight (J) hooks on the catch rates and numbers of white marlin and blue marlin released alive by the U.S. pelagic longline fleet in the Gulf of Mexico. North American Journal of Fisheries Management 28, 500–506. 10.1577/m07-089.1

Epperly SP, Watson JW, Foster DG, Shah AK (2012) Anatomical hooking location and condition of animals captured with pelagic longlines: The grand banks experiments 2002-2003. Bulletin of Marine Science 88, 513–527.

Fernandez-Carvalho J, Coelho R, Santos MN, Amorim S (2015) Effects of hook and bait in a tropical northeast Atlantic pelagic longline fishery: Part II—Target, bycatch and discard fishes. Fisheries Research 164, 312–321.

Foster DG, Epperly SP, Shah AK, Watson JW (2012) Evaluation of hook and bait type on the catch rates in the western North Atlantic Ocean pelagic longline fishery. Bulletin of Marine Science 88, 529–545.

Gabry J, Češnovar R (2021) cmdstanr: R Interface to “CmdStan.” Available at https://mc-stan.org/cmdstanr/reference/cmdstanr-package.html

Gallagher AJ, Orbesen ES, Hammerschlag N, Serafy JE (2014) Vulnerability of oceanic sharks as pelagic longline bycatch. Global Ecology and Conservation 1, 50–59.

Gilman E, Chaloupka M, Bach P, Fennell H, Hall M, Musyl M, Piovano S, Poisson F, Song L (2020) Effect of pelagic longline bait type on species selectivity: a global synthesis of evidence. Reviews in Fish Biology and Fisheries 30, 535–551. 10.1007/S11160-020-09612-0/TABLES/2

Gilman E, Chaloupka M, Swimmer Y, Piovano S (2016) A cross-taxa assessment of pelagic longline by-catch mitigation measures: conflicts and mutual benefits to elasmobranchs. Fish and Fisheries 17, 748–784. 10.1111/faf.12143

Gilman E, Kobayashi D, Swenarton T, Brothers N, Dalzell P, Kinan-Kelly I (2007) Reducing sea turtle interactions in the Hawaii-based longline swordfish fishery. Biological Conservation 139, 19–28. 10.1016/j.biocon.2007.06.002

Gilman E, Zollett E, Beverly S, Nakano H, Davis K, Shiode D, Dalzell P, Kinan I (2006) Reducing sea turtle by-catch in pelagic longline fisheries. Fish and Fisheries 7, 2–23. 10.1111/j.1467-2979.2006.00196.x

Godin AC, Carlson JK, Burgener V (2012) The effect of circle hooks on shark catchability and at-vessel mortality rates in longlines fisheries. Bulletin of Marine Science 88, 469–483. 10.5343/bms.2011.1054

Hall MA, Alverson DL, Metuzals KI (2000) By-catch: problems and solutions. Marine Pollution Bulletin 41, 204–219. 10.1016/S0025-326X(00)00111-9

Hiraoka Y, Kanaiwa M, Ohshimo S, Takahashi N, Kai M, Yokawa K (2016) Relative abundance trend of the blue shark *Prionace glauca* based on Japanese distant-water and offshore longliner activity in the North Pacific. Fisheries Science 82, 687–699. 10.1007/s12562-016-1007-7

Horodysky AZ, Graves JE (2005) Application of pop-up satellite archival tag technology to estimate postrelease survival of white marlin (*Tetrapturus albidus*) caught on circle and straight-shank (“J”) hooks in the western North Atlantic recreational fishery. Fishery Bulletin 103, 84.

Huang HW, Swimmer Y, Bigelow K, Gutierrez A, Foster DG (2016) Influence of hook type on catch of commercial and bycatch species in an Atlantic tuna fishery. Marine Policy 65, 68–75. 10.1016/j.marpol.2015.12.016

ICCAT (2019) Report of the 2019 Shortfin mako shark stock assessment update meeting. *Collective Volume of Scientific Papers*, ICCAT 76, 1–77.

Kay M (2023) tidybayes: Tidy data and geoms for Bayesian models. Available at 10.5281/zenodo.1308151

Kerstetter DW, Graves JE. (2006) Effects of circle versus J-style hooks on target and non-target species in a pelagic longline fishery. Fisheries Research 80, 239–250. 10.1016/j.fishres.2006.03.032

Kerstetter DW, Luckhurst BE, Prince ED, Graves JE (2003) Use of pop-up satellite archival tags to demonstrate survival of blue marlin (*Makaira nigricans*) released from pelagic longline gear. Fishery Bulletin 101, 939–941.

Kiyota M, Yokota K, Nobetsu T, Minami H, Nakano H (2004) Assessment of mitigation measures to reduce interactions between sea turtles and longline fishery. Proceedings of the International Symposium on SEASTAR 2000 and Bio-Logging Science (The 5th SEASTAR 2000 Workshop), 24–29.

Kruschke JK (2015) Doing bayesian data analysis: a tutorial with R, JAGS, and Stan. 2nd eds. (Academic Press, New York, USA)

Maunder MN, Punt AE (2004) Standardizing catch and effort data: a review of recent approaches. Fisheries Research 70, 141–159. 10.1016/j.fishres.2004.08.002

Melvin EF, Guy TJ, Read LB (2014) Best practice seabird bycatch mitigation for pelagic longline fisheries targeting tuna and related species. Fisheries Research 149, 5–18. 10.1016/j.fishres.2013.07.012

Pacheco JC, Kerstetter DW, Hazin FH, Hazin H, Segundo RSSL, Graves JE, Carvalho F, Travassos PE (2011) A comparison of circle hook and J hook performance in a western equatorial Atlantic Ocean pelagic longline fishery. Fisheries Research 107, 39–45. 10.1016/j.fishres.2010.10.003

Parga ML, Pons M, Andraka S, Rendón L, Mituhasi T, Hall M, Pacheco L, Segura A, Osmond M, Vogel N (2015) Hooking locations in sea turtles incidentally captured by artisanal longline fisheries in the Eastern Pacific Ocean. Fisheries Research 164, 231–237. 10.1016/j.fishres.2014.11.012

Pebesma E (2018) Simple features for R: Standardized support for spatial vector data. The R Journal 10, 439–446. 10.32614/RJ-2018-009

R Core Team (2023) R: A Language and Environment for Statistical Computing. Available at https://www.r-project.org/

Reinhardt JF, Weaver J, Latham PJ, Dell’Apa A, Serafy JE, Browder JA, Christman M, Foster DG, Blankinship DR (2018) Catch rate and at-vessel mortality of circle hooks versus J-hooks in pelagic longline fisheries: A global meta-analysis. Fish and Fisheries 19, 413–430. 10.1111/faf.12260

Santos MN, Coelho R, Fernandez-Carvalho J, Amorim S (2012) Effects of Hook and bait on sea turtle catches in an equatorial Atlantic pelagic longline fishery. Bulletin of Marine Science 88, 683–701. 10.5343/bms.2011.1065

Santos CC, Rosa D, Coelho R (2020) Progress on a meta-analysis for comparing hook, bait and leader effects on target, bycatch and vulnerable fauna interactions. Collective Volume of Scientific Papers ICCAT 77, 182–217.

Sims DW, Mucientes G, Queiroz N (2018) Shortfin mako sharks threatened by inaction. Science 359, 1342. 10.1126/science.aat0315

Skomal G, Bernal D (2010) Physiological responses to stress in sharks. In ‘Sharks and Their Relatives II.’ (Eds JC Carrier, JA Musick, MR Heithaus), pp. 475–506. (CRC Press, Boca Raton, USA)

Stokes L, Hataway D, Epperly S, Shah A, Bergmann C, Watson J, Higgins B (2011) Hook ingestion rates in loggerhead sea turtles *Caretta caretta* as a function of animal size, hook size, and bait. Endangered Species Research 14, 1–11. 10.3354/esr00339

Swimmer Y, Gutierrez A, Bigelow K, Barceló C, Schroeder B, Keene K, Shattenkirk K, Foster DG (2017) Sea turtle bycatch mitigation in U.S. longline fisheries. Frontiers in Marine Science 4, 260. 10.3389/FMARS.2017.00260

Stan Development Team (2021) Stan modeling language users guide and reference manual, 2.28.2. Available at https://mc-stan.org

Wallace BP, Kot CY, Dimatteo AD, Lee T, Crowder LB, Lewison RL (2013) Impacts of fisheries bycatch on marine turtle populations worldwide: toward conservation and research priorities. Ecosphere 4, 1–49. 10.1890/ES12-00388.1

Watson JW, Epperly SP, Shah AK, Foster DG (2005) Fishing methods to reduce sea turtle mortality associated with pelagic longlines. Canadian Journal of Fisheries and Aquatic Sciences 62, 965–981. 10.1139/f05-004

Wickham H, Averick M, Bryan J, Chang W, McGowan LD, François R, Grolemund G, Hayes A, Henry L, Hester J et al. (2019) Welcome to the tidyverse. Journal of Open Source Software 4, 1686. 10.21105/joss.01686

Yokota K, Kiyota M, Minami H (2006a) Shark catch in a pelagic longline fishery: Comparison of circle and tuna hooks. Fisheries Research 81, 337–341. 10.1016/j.fishres.2006.08.006

Yokota K, Kiyota M, Okamura H (2009) Effect of bait species and color on sea turtle bycatch and fish catch in a pelagic longline fishery. Fisheries Research 97, 53–58. 10.1016/J.FISHRES.2009.01.003

Yokota K, Minami H, Kiyota M (2006b) Measurement-points examination of circle hooks for pelagic longline fishery to evaluate effects of hook design. Bulletin of Fishery Research Agency 17, 83–102

Yokota K, Minami H, Kiyota M (2011) Effectiveness of tori-lines for further reduction of incidental catch of seabirds in pelagic longline fisheries. Fisheries Science 77, 479–485. 10.1007/s12562-011-0357-4

